# A classification of structured coalescent processes with migration, conditional on the population pedigree

**DOI:** 10.64898/2026.02.18.706396

**Authors:** Sabin Lessard, Terry Easlick, John Wakeley

## Abstract

Recent analyses of the effects that organismal genealogies or pedigrees of populations have on times to common ancestry for samples of genetic data are extended to cases of population subdivision and migration. Traditional coalescent models marginalize over pedigrees. A finding of a ‘pedigree effect’ implies that data analysis and interpretation should not be based on the corresponding traditional coalescent model but rather on a coalescent model obtained by conditioning on the pedigree. We apply a straightforward test based on the distribution of pairwise coalescence times to four previously described scenarios of subdivision and migration. These scenarios are defined by the relative magnitudes of four parameters: the number of the local populations or demes, the deme size, the migration fraction, and the probability that migration can occur at all. We find pedigree effects in three scenarios. In two, the effect is weak if the deme size is large. The one scenario without a pedigree effect corresponds to the well known structured-coalescent model. The one scenario with a persistent pedigree effect even in the limit as the deme size tends to infinity involves long periods without gene flow interrupted by pulses of migration. We illustrate our results using simulations and numerical analysis.

## Introduction

Coalescent models describe genetic ancestry in order to predict genetic variation in a sample from a population. These continuous-time models measure time backwards from the present to the past and make minimal reference to the population beyond the sample. Generally, they are obtained by taking the limit as the population size tends to infinity with time rescaled so the coalescence rate for a pair of ancestral lineages is equal to one. Their development can be traced through key papers such as Ewens (1972) and Watterson (1975), and considered mature in Kingman (1982a,b,c), Hudson (1983a,b) and Tajima (1983). After these initial descriptions for well-mixed populations of constant size without selection, coalescent models were extended to cover various kinds of population structure, changes in population size, and selection; see the detailed overview by Nordborg (2001) or fuller treatments by Hein et al. (2005), Berestycki (2009) and Wakeley (2009).

When applied to diploid biparental organisms, these traditional coalescent models describe gene genealogies or other aspects of genetic ancestry without reference to the organismal genealogy or pedigree of the population. They instead describe ancestral genetic processes marginal to the pedigree. Their predictions are averages over all possible pedigrees. See Diamantidis et al. (2024) for a review of this practice and its historical development. Ball et al. (1990) were the first to ask whether distributions of gene genealogies conditional on the population pedigree might differ substantially from the marginal distributions. The motivation behind this question is that there is an actual pedigree of any population, and it is the structure of this pedigree that determines possible genetic ancestries of a sample. If the conditional and marginal distributions of gene genealogies differ, the utility of the corresponding traditional coalescent model may be limited.

Distributions of coalescence times or other aspects of genetic ancestry conditional on the population pedigree are predictions about what should be observed in data from a very large number of unlinked loci in a sample of individuals. This is the sampling structure implicit in many applications of traditional coalescent models, for example when likelihoods are multiplied across loci in a sample of individuals using programs such as *∂a∂i* (Gutenkunst et al., 2009), *momi*2 (Kamm et al., 2020) and *fastsimcoal*2 (Excoffier et al., 2013, 2021), or when sample site-frequency spectra are compared to model predictions (Braverman et al., 1995; Tajima, 1989; Fu, 1995; Achaz, 2009; Gao and Keinan, 2016; Ragsdale et al., 2018; Patel et al., 2025). However, it is only conditional on the pedigree that patterns of variation at unlinked loci are independent.

We note that sometimes the predictions used in these applications come from forward-time Wright-Fisher diffusion theory rather than backward-time coalescent models (Ewens, 2004). But predictions about site frequencies or other measures of genetic variation from diffusion theory are also marginal to the population pedigree. They are derived by averaging over the outcomes of reproduction. Traditional coalescent models are backward-time versions, or dual processes, of the corresponding forward-time diffusion models (Donnelly, 1986; Ewens, 1990; Möhle, 1999).

Ball et al. (1990) showed using simulations that the pedigree has little effect if the population is well mixed and of constant size without selection, provided the population size is reasonably large (500 individuals). Subsequent simulations over a large range of population sizes upheld this conclusion (Wakeley et al., 2012), in particular that distributions of coalescence times among unlinked loci in a sample of individuals conditional on their shared pedigree conform well to predictions from the standard neutral coalescent model of Kingman (1982a,b,c), Hudson (1983a,b) and Tajima (1983). Tyukin (2015) proved this for a monoecious diploid population meeting the assumptions of this model: in the limit as the population size tends to infinity, conditioning on the pedigree or marginalizing over it leads to the same result which is the standard neutral coalescent model.

This type of robustness to the pedigree cannot be assumed to hold for populations outside the domain of the standard neutral coalescent model. For both modeling and inference, it is important to know for all kinds of populations: Does the limiting coalescent process conditional on the population pedigree differ from the corresponding traditional coalescent model? A negative answer, that there is no pedigree effect, means we can keep using the statistical models and methods of inference developed for the corresponding traditional coalescent model. A positive answer means we need to develop new models and methods. Here, we provide answers for different types of subdivided populations comprised of discrete local subpopulations, or demes, connected by migration.

We do this for an extended version of Wright’s finite island model (Wright, 1931, 1943; Maruyama, 1970; Latter, 1973) in which the times and sizes for migration events can be decoupled, as detailed in Structured population model with gametic migration. Specifically, in Four kinds of conditional coalescent models we consider four limits of the model under different assumptions about the relative magnitudes of its parameters. We identify cases in which the conditional coalescent process and the corresponding pedigree-averaged process differ and cases in which they are the same. Briefly and before getting into the technical details, these limits and our findings are as follows.

The first limit is the one which gives the well known structured coalescent model in the pedigree-averaged case (Takahata, 1988; Notohara, 1990; Herbots, 1997; Wilkinson-Herbots, 1998). This is a model for large demes and correspondingly small migration probabilities. Of the four models we consider, this is by far the most frequently applied (Nordborg, 1997; Beerli and Felsenstein, 2001; Pinho and Hey, 2010; Lohse et al., 2011, 2016; Costa and Wilkinson-Herbots, 2017; Müller et al., 2017; Beerli et al., 2019). In this case, our results suggest that coalescence conditional on the pedigree can be described using the available structured coalescent model.

The second limit is the many-demes limit (Wakeley, 1998, 2003; Wakeley and Takahashi, 2004) of the island migration model (Wright, 1931, 1943; Maruyama, 1970; Latter, 1973). This is a model for finely subdivided populations or metapopulations (Pannell and Charlesworth, 2000; Pannell, 2003; Hanski and Gaggiotti, 2004). In this case, we show that the conditional and pedigree-averaged pairwise coalescent processes differ, but only for ancestral lines starting in the same deme. An essentially similar result has recently been shown to hold for populations with appreciable levels of self-fertilization (Newman et al., 2025; Fan et al., 2025). Roughly speaking, and as elaborated in Lessard and Wakeley (2004) in the context of a traditional coalescent model, demes play the role of individuals and migration plays the role of outcrossing.

The third limit is the low-migration limit of Lande (1979, 1985) and Slatkin (1981). This is a separation of time scales approximation in which events within finite-sized demes resolve quickly against a backdrop of long waiting times between migration events, due to a constant low probability of migration. It has been used to understand forward-time dynamics of frequency-dependent selection (Lande, 1979, 1985; Slatkin, 1981; Cherry, 2003; Nishino and Tajima, 2004, 2005; Ladret and Lessard, 2007) as well as to describe patterns of coalescence and neutral genetic variation (Takahata, 1991; Notohara, 2001; Baronchelli and Pastor-Satorras, 2009; Burden and Griffiths, 2018). In this case, we show that the conditional and pedigree-averaged coalescent processes differ for ancestral lines starting in different demes.

The fourth limit is a model of stochastic migration (Sved and Latter, 1977; Nagylaki, 1979; Latter and Sved, 1981; Whitlock, 1992) in which a potentially large fraction of a deme is replaced by migrants but these events occur rarely, at random times and with no gene flow in between (Peniston et al., 2024). Our model is similar to the ones which Gaggiotti and Smouse (1996), Rice and Papadopoulos (2009), Yamaguchi and Iwasa (2013, 2017) and Aubree et al. (2022) studied in other contexts. We refer to this as the rare-migration limit. In this final case as well, we show that the conditional and pedigree-averaged coalescent processes differ as in the low-migration limit.

The three models for which we do find a pedigree effect have in common that the deme size is fixed and finite in the respective limit, though we do note that the total population size becomes infinite in the many-demes limit. Given that single finite populations shows pedigree effects (Wakeley et al., 2012; Tyukin, 2015), we further ask whether our results for these three models depend on the assumption of finite deme size. In the many-demes limit and the low-migration limit, the effect of conditioning on the pedigree becomes negligible as the deme size increases. In the rare-migration limit, the pedigree effect persists even as the deme size tends to infinity.

These will be our main findings. They primarily support the continued use of the widely used structured coalescent model of population subdivision and migration with demes of large enough size, which was never justified with respect to the population pedigree. They also point toward the development of coalescent models conditional on the population pedigree for application to metapopulations, populations with very limited gene flow, and populations with large migration events or demes of small size. In addition, in the next section we provide a new explanation of a method for assessing whether conditioning on the pedigree leads to a different coalescent model than averaging over the pedigree or whether both approaches give the same results in the limit.

Our results complement a growing body of work on coalescent processes conditional on the population pedigree. Tyukin (2015) found no pedigree effect in the usual coalescent limit for a single population meeting the assumptions of the standard neutral coalescent model. Pedigree effects have been found for populations in which some individuals can have very large numbers of offspring (Diamantidis et al., 2024; Alberti et al., 2025) and populations in which there is an appreciable chance of self-fertilization (Newman et al., 2025; Fan et al., 2025). Our results for the structured-coalescent limit and the rare-migration limit overlap with the recent results of Newman (2026), who studied the conditional coalescent processes of samples of arbitrary size in a general model of subdivision and migration in the limit of a large deme size with the possibility of large reproduction events.

### The conditional survival function and its variance

In this work we characterize distributions of single-locus pairwise coalescence times, conditional on the population pedigree and on how the two genes are sampled in the current generation. In general, we will be concerned with two types of samples: two genes at a locus in a single individual chosen at random from the population, and two genes in two individuals sampled from two different demes. Our methods will cover other sampling configurations as well. For illustration, in this section we focus on the two genes within an individual.

In addition, we consider the most familiar way that coalescence times are defined for diploids, namely in the limit as the population size *N* tends to infinity with time measured in units of 2*N* generations. We denote the coalescence time by *τ*^(*N*)^, keeping the dependence on *N* explicit. We use 𝒜 to represent the population pedigree, leaving the dependence on *N* implicit to simplify the notation. We assume that 𝒜 is the outcome of a random process of reproduction (as in Structured population model with gametic migration) and that it includes the information of which individual was sampled, so that the conditional coalescence time *τ*^(*N*)^ given 𝒜 is well defined.

Following Diamantidis et al. (2024), we characterize the distribution of the coalescence time using the conditional survival function

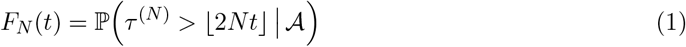

for *t* ≥ 0, where ⌊2*Nt*⌋ designates the largest integer less than or equal to 2*Nt*. Thus *t* is time measured in units of 2*N* generations, and (1) is the probability that the two ancestral lines starting from the sampled individual coalesce more than ⌊2*Nt*⌋ generations ago, given the population pedigree. Note that 1 − *F*_*N*_ (*t*) is the cumulative distribution function of the coalescence time.

Our goal is to describe the properties of *F*_*N*_ (*t*) as *N* → ∞. In particular, we wish to know whether it retains its dependence on the random population pedigree in the limit or it becomes a deterministic function only of *t* as in traditional coalescent models. By the tower property of conditional expectation, we have

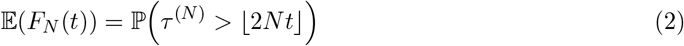

for *t* ≥ 0 as the mean of the survival function over all possible population pedigrees. The limit of (2) as *N* → ∞ is how the coalescence time would be characterized under the traditional approach of coalescence theory. Now, if Var(*F*_*N*_ (*t*)) tends to zero as *N* → ∞, then *F*_*N*_ (*t*) converges in mean square, and therefore in distribution, to its mean (2) for every fixed *t* ≥ 0. This implies that the limiting conditional coalescent model must coincide with the limiting unconditional one. But if Var(*F*_*N*_ (*t*)) does not tend to zero, then *F*_*N*_ (*t*) remains a stochastic function, depending on the pedigree, even as *N* → ∞.

The variance of *F*_*N*_ (*t*) can be computed by considering the coalescence times 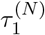 and 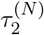 of the ancestral lines that start from two pairs of genes at two unlinked loci in the same individual chosen at random, which are conditionally independent and identically distributed given the population pedigree. Again following Diamantidis et al. (2024), we have the identities

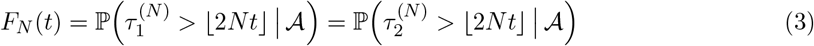

and

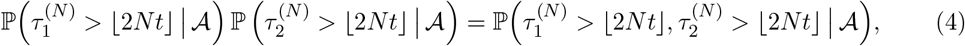

from which

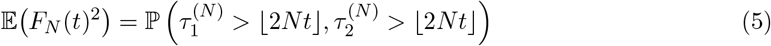

for *t* ≥ 0. In (2), (4) and (5) we have what we need to compute Var(*F*_*N*_ (*t*)).

In Four kinds of conditional coalescent models, we apply this logic to four different limits of the population model presented in Structured population model with gametic migration, computing the mean and the variance of the survival function in each case. For each model, we ask whether the effect of the pedigree persists in the relevant limit. In fact, only one of these four models has *N* → ∞. The other limits concern aspects of the migration process or the number of demes.

Note that (2) and (5) are averages over the population pedigree. Conveniently, there is no need to specify the whole pedigree or to compute the conditional survival function (1). For any population model which can be described as a Markov chain, (2) and (5) can be computed from the transition matrix. It is not even necessary to specify all aspects of the pedigree process (i.e. all details of reproductive outcomes among individuals) in the relevant Markov chain, insofar as these do not affect the averages (2) and (5). In what follows, we take advantage of this by assuming that reproduction occurs among monoecious individuals via random union of gametes from an infinite gamete pool which may include some proportion of migrant gametes. Describing the model in this way minimizes the number of states in the Markov chains required to compute (2) and (5). Some key extensions are treated in Robustness to diploid migration and dioecy and the Supplementary Material Sections 4 and 5. Extensions to samples larger than two are taken up in the Discussion.

The method of assessing convergence using (2) and (5) was developed to analyze random walks in random environments (Bolthausen and Sznitman, 2002; Birkner et al., 2013). In coalescent models, the two conditionally independent processes supposed in the method correspond exactly to the Mendelian transmission of two unlinked loci (having coalescence times 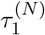 and 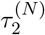. So, the computations involved are directly relevant to the genetic ancestries of unlinked loci. The statement that Var(*F*_*N*_ (*t*)) → 0 is the same as the statement that 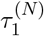 and 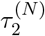 retain their independence in the limit even when averaging over pedigrees as in (5). On the other hand, if *F*_*N*_ (*t*) remains a stochastic function of the pedigree, it means that 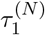 and 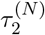 become non-independent in the limit due to the random pedigree they share.

### Structured population model with gametic migration

Our goal is to identify types of structured populations for which the limiting coalescent process depends on the population pedigree. For simplicity, we adopt a symmetric island model in which *D* demes each containing *N* diploid monoecious individuals are all equally connected by migration, and reproduction occurs via an effectively infinite gamete pool. Such a model would ordinarily include a single migration parameter, but here we will have two. The usual single parameter, which we also include, is denoted *m*. This is the proportion of each deme’s gamete pool that comes from the other demes. The second parameter controls the timing of migration events across the entire population. Specifically, we define *α* as the probability that any migration can occur at all in a generation. With probability 1 − *α*, no migration is possible, so each deme’s gametes come from within that deme. With probability *α*, all demes’ gamete pools include proportions 1 − *m* and *m* of non-migrant and migrant gametes, respectively. Migration being symmetric means that fractions *m/*(*D* − 1) come from each of the other *D* − 1 demes.

We calculate the mean and the variance of the conditional distribution function of pairwise coalescence times. This entails describing backward-time Markov processes allowing (2) and (5) to be computed. Under our simple model, the average (2) can be computed by tracking just three gametic states in the ancestral process. Our starting sample includes the pair of genes at a single locus in an individual chosen at random in the current generation. These are the two genes in the two gametes which united to form the individual. In the model, they are no different than two genes in two gametes which went to different individuals in the same deme. Then it suffices to consider just three possible states of the ancestral lines of the two genes in the sample: [•] for one in one deme, i.e. the coalescent state, [••] for two in the same deme, and [•][•] for two in two different demes. Throughout, we use square brackets in this way to delineate demes. When necessary, we use parentheses to delineate individuals, so for example either [(••)] or [(•)(•)] for two gametes in the same deme but in the same individual or in different individuals, respectively.

The corresponding transition probability matrix for the ancestral process of a sample of two genes at a single locus is

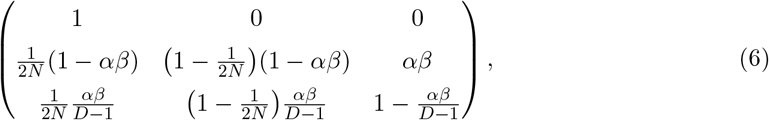

in which

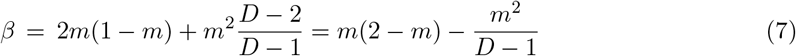

represents the conditional probability for two gametes chosen at random in the same deme to come from two different demes given that migration occurs. Note that the conditional probability for two gametes chosen at random in different demes to come from the same deme is then given by *β/*(*D* − 1). In the next section, Four kinds of conditional coalescent models, we use (6) to compute the mean of the conditional survival function (2) and describe how two-locus gametic states can be tracked in order to compute (5).

### Four kinds of conditional coalescent models

Consider a monoecious diploid population subdivided into *D* demes of size *N* and undergoing discrete, nonoverlapping generations. From one generation to the next, gametes are produced in equal large numbers by the individuals in all the demes. With probability *α*, a fraction 1 − *m* of gametes stay in the deme where they were produced while a fraction *m* migrate uniformly to the other demes. Then, *m* is the fraction of gametes in each deme that come from the other demes and *m/*(*D* − 1) the fraction that come from each one. Otherwise, with probability 1 − *α*, all gametes stay in the deme where they were produced. Random sampling of *N* pairs of gametes within each deme starts the next generation.

We are interested in the coalescence time of ancestral lines that start in the current generation from a pair of genes at a single locus in an individual chosen at random in the population or in two different individuals chosen at random in different demes. We consider these coalescence times under four assumptions: (1) *N* → ∞ with *Nm* fixed for *D* = 2 and *α* = 1 (the structured-coalescent limit); (2) *D* → ∞ with *N* and *m* fixed for *α* = 1 (the many-demes limit); (3) *m* → 0 with *N* fixed for *D* = 2 and *α* = 1 (the low-migration limit); and (4) *α* → 0 with *N* and *m* fixed for *D* = 2 (the rare-migration limit).

In case (1), the coalescence time, rescaled so that it is measured in units of 2*N* generations, is represented by *τ*^(*N*)^. Its conditional survival function given the population pedigree is *F*_*N*_ (*t*) for *t* ≥ 0, as defined in The conditional survival function and its variance. The first two moments of the conditional survival function at rescaled time *t* ≥ 0, namely, 𝔼(*F*_*N*_ (*t*)) and 𝔼 *F*_*N*_ (*t*)^2^, will be obtained in the limit by considering the transition probabilities from one generation to the previous one for possible states of the ancestral lines (in the same deme or in different demes, in the same individual or in different individuals). The first moment corresponds to the probability that there are still two ancestral lines at time *t* starting from the pair of genes at time 0. The second moment corresponds to the probability that there are still four ancestral lines at time *t* starting from two pairs of genes at two unlinked loci at time 0.

In cases (2), (3) and (4), the parameter *N* will be replaced by *D, m*, and *α*, respectively, and subsequently the unit of time 2*N* generations will be replaced by *D*, 1*/m*, and 1*/α* generations, respectively. For ease of comparison with the known results of this well-studied model, we choose 2*N* rather than *N* for the time scale in the structured coalescent case (1), and similarly define the parameter *M* = 4*Nm* as in Wilkinson-Herbots (1998) Section 4. To illustrate the key principles, we restrict ourselves to simplified versions of the general model in each case. In Supplementary Material, we treat a number of familiar extensions of these simplified models.

### Structured-coalescent limit

In this subsection, we assume *D* = 2 demes of size *N* and migration fraction *m* = *M/*(4*N*) with probability *α* = 1 in each generation. Starting from a pair of genes at the same locus in an individual chosen at random in a deme chosen at random, the ancestral lines backward in time can be in three possible states ordered as follows: [•] for only one in one deme, [••] for two in the same deme, and [•][•] for two in two demes. Our assumptions of random union of gametes and gametic migration allow for this reduced state space, in that we do not need to specify whether two ancestral lines in the same deme, [••], are in the same individual or in different individuals.

The transition probabilities from one generation to the previous one are given by a transition matrix that can be written in the form

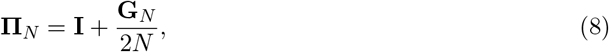

where **I** is an identity matrix and

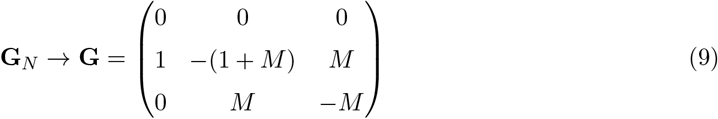

as *N* → ∞. By the definition of the matrix exponential (see Möhle (1998a) for a proof), we have

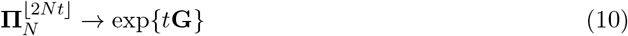

as *N* → ∞. Diagonalizing the matrix **G** leads to

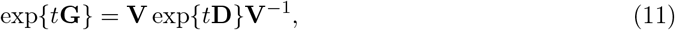

where

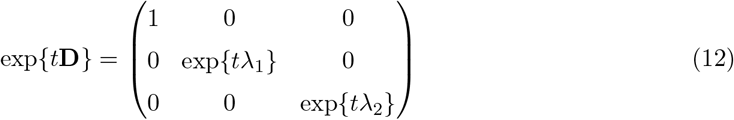

and

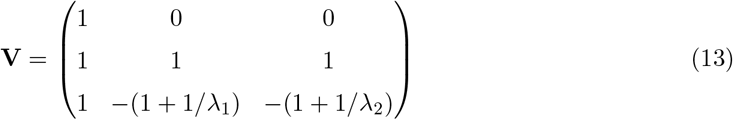

with

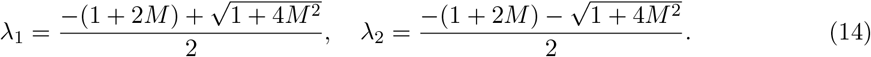

Using the above convergence result, the limit of

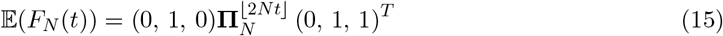

as *N* → ∞ is given by

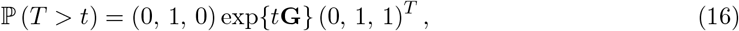

where the random variable *T* represents the time for a continuous-time Markov chain whose rate matrix is **G** to fix in state [•] starting from state [••]. Note that this rate matrix corresponds to the situation where two ancestral lines in the same deme coalesce at rate 1 while each one migrates from one deme to the other at rate *M/*2, all these events occurring independently.

Next we consider the possible states for the ancestral lines of two pairs of genes at two unlinked loci, 1 and 2, in the same individual. They are ordered as follows, in which •, ○, and 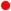 represent gametes ancestral at locus 1 only, at locus 2 only, and at both loci, respectively, and brackets delineate demes:

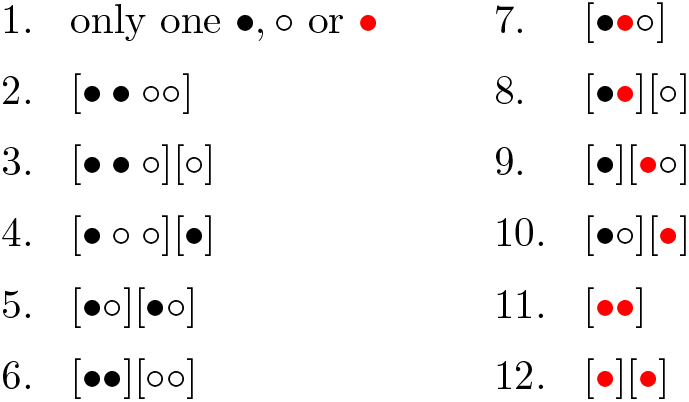

Because our focus is on the survival function, we lump all coalescent events into state 1.

The transition matrix from one generation to the previous one takes the form

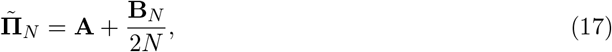

where

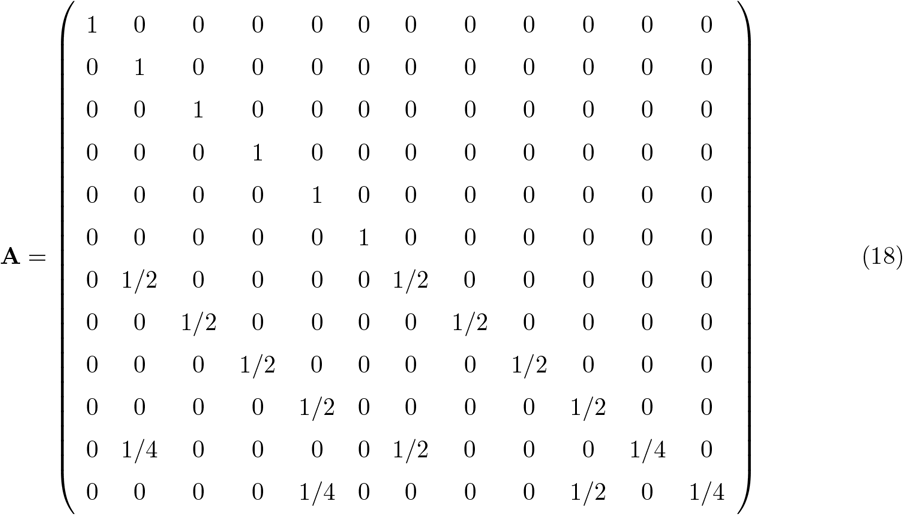

and **B**_*N*_ → **B** as *N* → ∞ with the first 6 rows of **B** given by

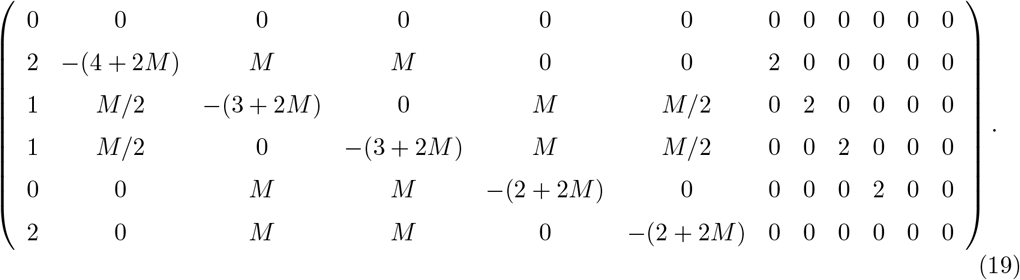

Lemma 1 of Möhle (1998a) guarantees that

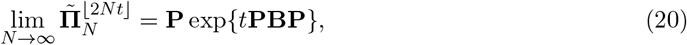

where

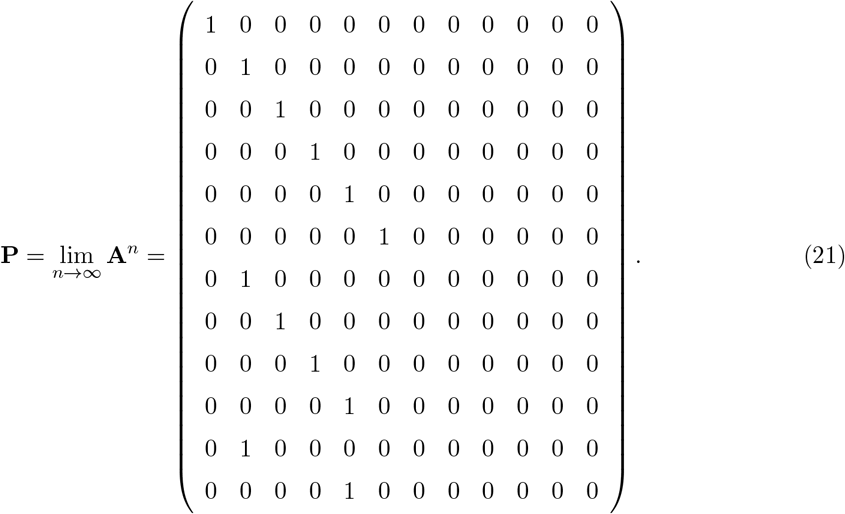

This implies that, in the limit as *N* → ∞, there are instantaneous transitions to states in the set *S*_1_ = {1, 2, …, 6} if the initial state belongs to the set {7, 8, …, 12}, while once in the state space *S*_1_ we have a continuous-time Markov chain whose rate matrix 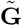 is given by the entries of **PBP** corresponding to the elements of *S*_1_, that is,

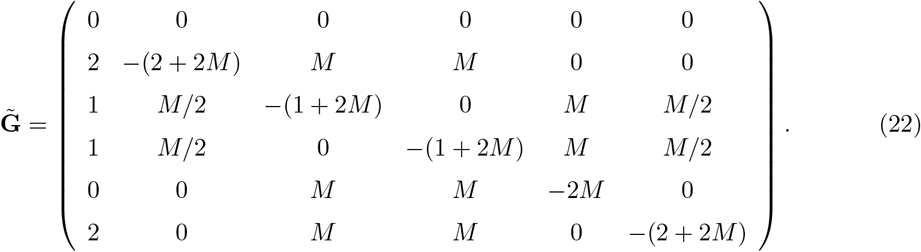

This is the rate matrix for the ancestral lines of two pairs of genes at two loci, 1 and 2, if the ancestral lines at each locus coalesce or migrate as previously and independently of the ancestral lines at the other locus. Therefore, we have

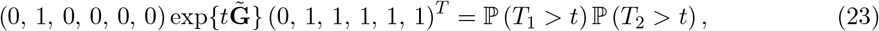

where *T*_1_ and *T*_2_ represent two independent random variables having the same probability distribution as *T* previously introduced. But this is the limit of

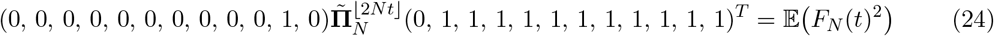

as *N* → ∞, since there is an instantaneous transition from state 11 to state 2 in the limit of a large deme size. We conclude that

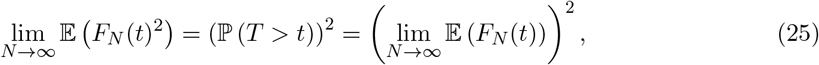

which implies that

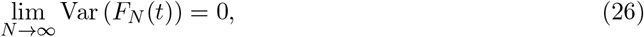

for *t* ≥ 0. The same conclusion can be drawn if the initial state is [•][•] instead of [••]. Conditioning on the pedigree and marginalizing over the pedigree give the same predictions about the distribution of pairwise coalescence times in the structured-coalescent limit.

#### Many-demes limit

Here we consider fixed *N* and *m*, with *D* → ∞. We set *α* = 1 for simplicity and for comparison with previous results from traditional many-demes coalescent models (Lessard and Wakeley, 2004; Wakeley, 2004a). We denote the possible states for the pair of ancestral lines as previously: [•] for only one in one deme, [••] for two in the same deme, and [•][•] for one in each of two different demes. Then, denoting (6) with *α* = 1 as **Π**_*D*_, we have

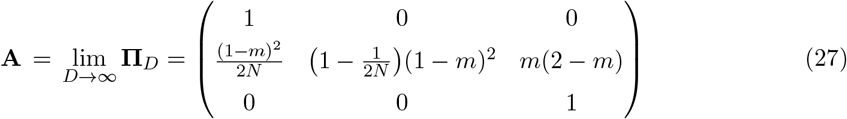

and

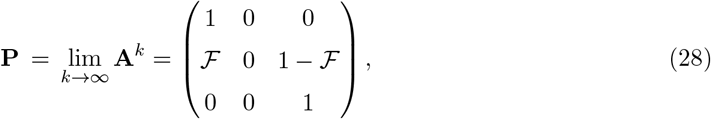

in which the coefficient

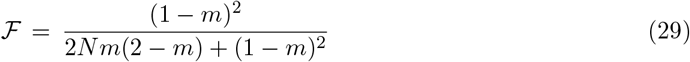

is equivalent to one definition of Wright’s *F*_*ST*_, namely the probability that the first event in the ancestry of two genetic lineages in the same deme is a coalescent event rather than a migration event (Wright, 1951; Slatkin, 1991; Rousset, 2002). Furthermore,

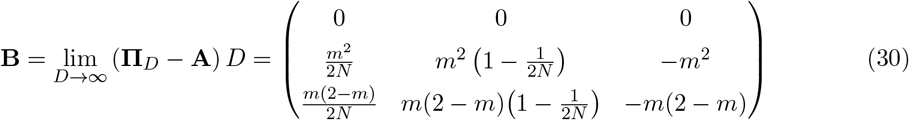

and

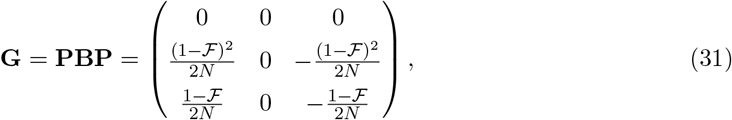

using the fact that

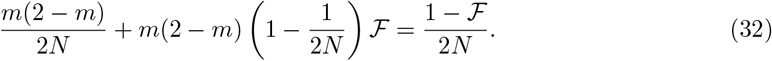

By Lemma 1 of Möhle (1998a), we have

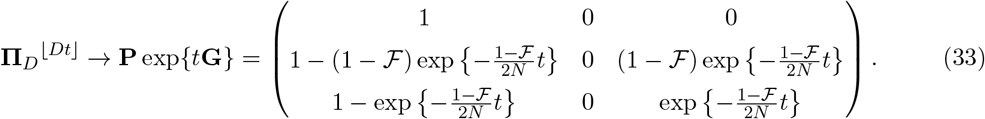

Therefore, the conditional survival function *F*_*D*_(*t*) = ℙ (*τ* ^(*D*)^ *>* ⌊*Dt*⌋ | 𝒜 satisfies

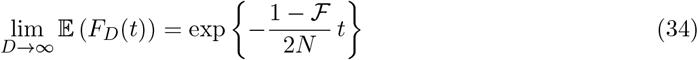

if the initial state is [•][•], but

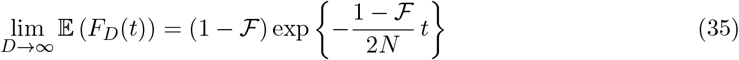

if the initial state is [••], for *t* ≥ 0. Thus, comparing (35) to (34), a sample in initial state [••] has probability ℱ of instantaneous coalescence, whereas with probability 1 − ℱ it follows the same coalescent process as a sample in initial state [•][•].

For two pairs of ancestral lines at two unlinked loci, we order the possible states as follows:

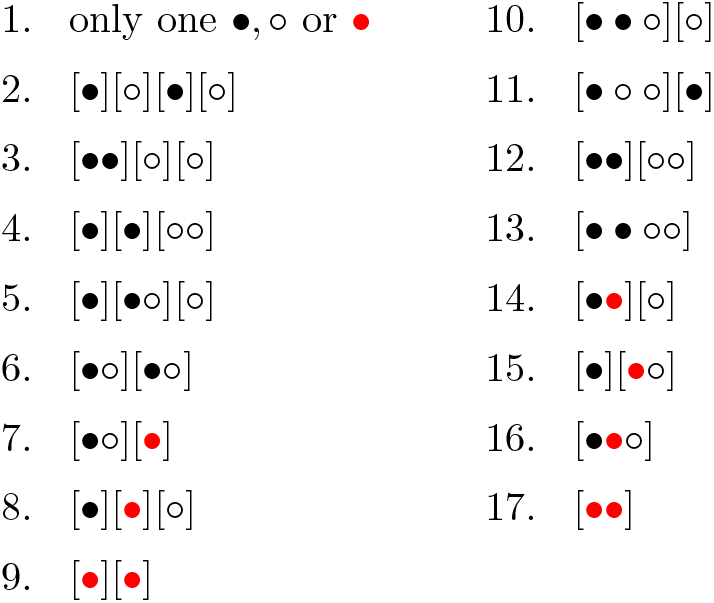

The transition matrix from one generation to the previous one can be expressed as

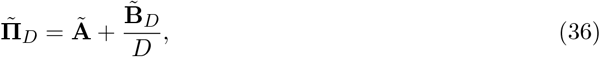

where **Ã** is the transition matrix in the limit of an infinite number of demes. In this limit, all states are transient but states 1 and 2, which are absorbing. Moreover, with respect to the subsets of states 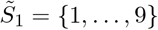 and 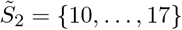, we have

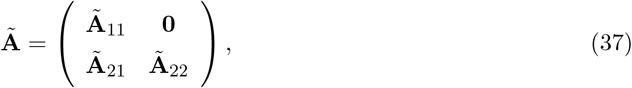

where

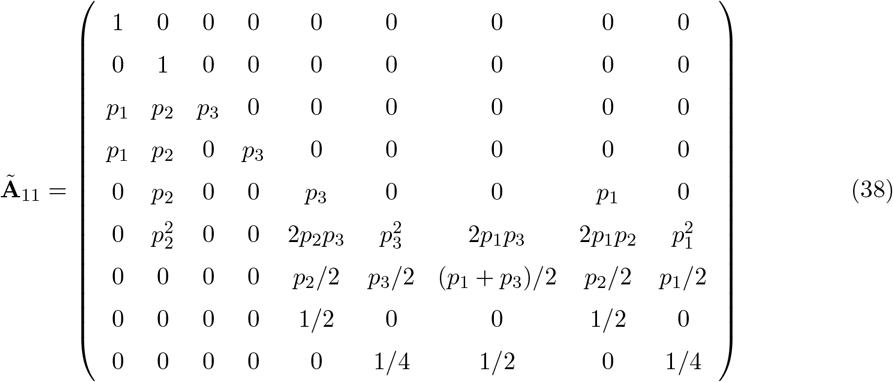

with

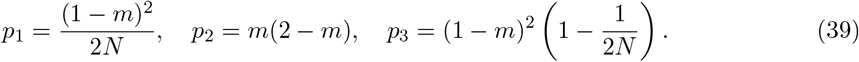

Therefore, the ergodic theorem for discrete-time Markov chains on a finite state space (see, e.g., Karlin and Taylor (1975)) ensures that

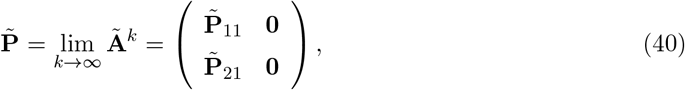

where

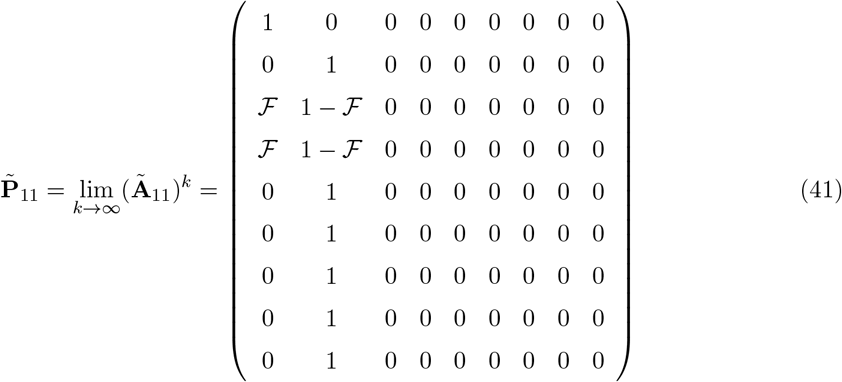

and

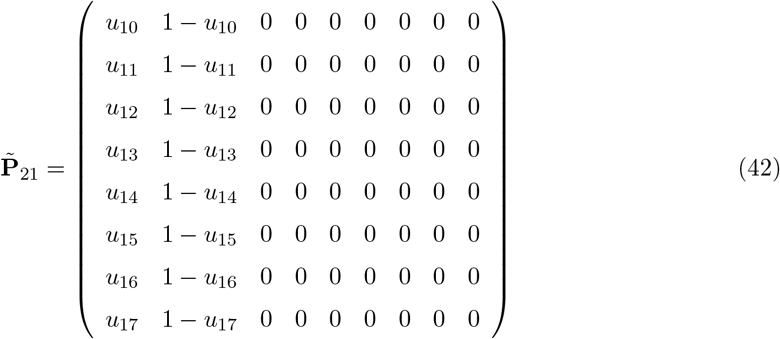

for some absorption probabilities *u*_10_, …, *u*_17_. In Supplementary Material Section 1.1 we find that *u*_10_ = *u*_11_ = *u*_14_ = *u*_15_ = ℱ and *u*_12_ = 1 −(1 −ℱ)^2^. These may be understood intuitively by noting that in the process specified by **Ã**: the outcomes for lineages in different demes are independent; two • lineages or two ○ lineages in the same deme will eventually coalesce with probability ℱ or disperse to different demes with probability 1 − ℱ; but two lineages of different types in the same deme must eventually disperse to different demes. The lengthy expressions for *u*_13_, *u*_16_ and *u*_17_, which satisfy equations (S7), (S8) and (S9), are shown in Supplementary Material Section 1.1.

On the other hand, the first two rows of

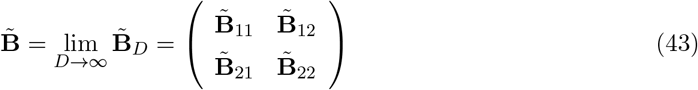

are given by

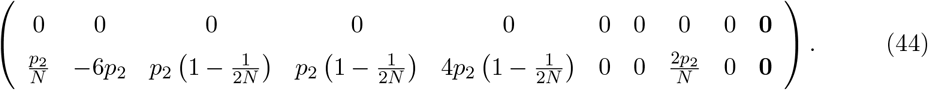

Lemma 1 of Möhle (1998a) ensures that

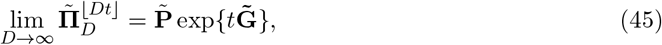

where

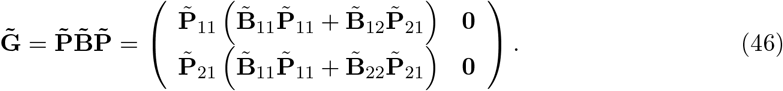

The first two rows of this matrix, which correspond to the first two rows of 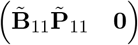, are

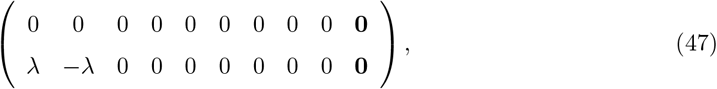

where

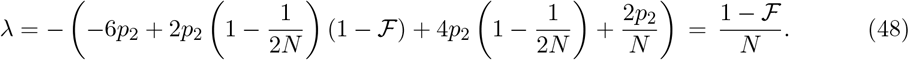

Therefore, in the limiting process with initial state 9, or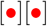, so that both pairs of lines start from the same chromosomes in two different demes, there is an instantaneous transition to the dispersed state 2, or [•][○][•][○], with all four lines in different demes and this is followed by a transition to the fixation state 1 at rate (1 − ℱ)*/N*. This implies that

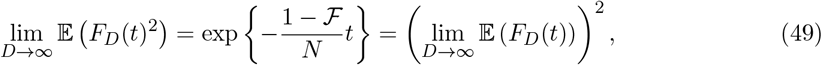

and then lim_*D*→∞_ Var (*F*_*D*_(*t*)) = 0, for *t* ≥ 0, if the initial state is [•][•].

With initial state 17, or 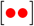, such that both pairs of lines start from the same chromosomes in the same deme, there will be an instantaneous transition to either the coalescent state 1 with probability *u*_17_ or to the dispersed state 2 with the complementary probability 1 − *u*_17_, from which the transition to the absorbing state 1 will occur at rate (1 − ℱ)*/N*. With *u*_17_ from (S7), (S8) and (S9) in the Supplementary Material Section 1.1, denoted 𝒰 for convenience, we conclude that

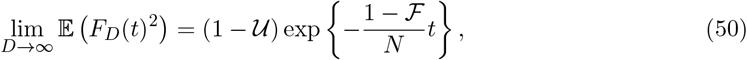

and then

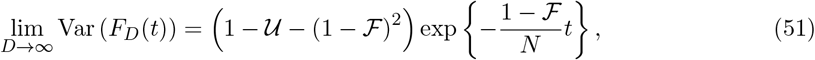

for *t* ≥ 0, if the initial state is [••]. The quantity 1 − 𝒰 − (1 − ℱ)^2^ depends on *N* and *m* and is complicated, but it is clearly not equal to zero (see Supplementary Material Sections 1.1 and 1.3). In this way, the many-demes coalescent process conditional on the population pedigree differs from the many-demes coalescent process marginal to the population pedigree.

So far in this section we have assumed that *N* and *m* are fixed as *D* → ∞. Our result (51) contrasts with the result (26) in the previous section Structured-coalescent limit where the effect of the population pedigree became negligible. However, that was in a different limit: *N* → ∞ with *m* = *M/*(4*N*). Previous, traditional many-demes coalescent models which implicitly average over the pedigree have often included *M* = 4*Nm* as a parameter, with comparatively little attention paid to finite *N* and fixed *m* (Lessard and Wakeley, 2004; Wakeley, 2004b).

Thus here we also ask whether the effect of the population pedigree in the many-demes limit becomes negligible if we further take *m* = *M/*(4*N*) and *N* large enough. As shown in Supplementary Material Section 1.2 where we express **Ã** in (36) as 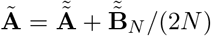 and then apply Lemma 1 of Möhle (1998a) as *N* → ∞, we have

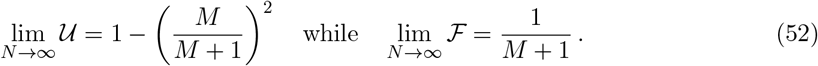

In other words, with *M* = 4*Nm* kept constant as *N* → ∞, the term 1 − 𝒰 − (1 − ℱ)^2^ in (51) tends to zero and the pedigree effect in the many-demes limit becomes negligible.

#### Low-migration limit

Here we consider *D* = 2 demes of fixed size *N* in the limit *m* → 0, with time rescaled accordingly and still with *α* = 1. The analysis follows the pattern of the previous two sections, Structured-coalescent limit and Many-demes limit, but we defer the details to Supplementary Material Section 2. Briefly, we define two state spaces and associated processes: for a single pair of ancestral lines (at one locus) and for two conditionally independent pairs of ancestral lines (at two unlinked loci). We write the transition matrix for the single-pair process as **Π**_*m*_ = **A** + *m***B**_*m*_, and the transition matrix for the two-pairs process as 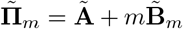, then apply Lemma 1 of Möhle (1998a) with time measured in units of 1*/m* generations.

Two ancestral lines in initial state [••] will coalesce immediately regardless of the pedigree, because opportunities for migration are vanishingly rare in this limit. However, computing the first two moments of the conditional survival function given an initial state [•][•], represented here by *F*_*m*_(*t*) = ℙ(*τ* ^(*m*)^ *>* ⌊*t/m*⌋ | 𝒜 and in the limit as *m* → 0, gives

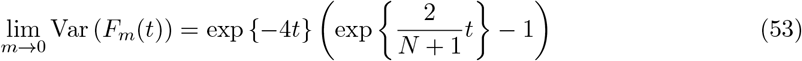

for *t* ≥ 0. Thus in the low-migration limit the coalescent process conditional on the population pedigree differs from the coalescent process marginal to the population pedigree.

Note that this limiting variance (53) vanishes as *N* → ∞. In Structured-coalescent limit, which had *N* → ∞ with *m* → 0 proportionally, the conditional and marginal coalescent processes were the same. In addition, in Many-demes limit, enforcing this same structured-coalescent limit on the end result (51) made the effect of the pedigree disappear. Similarly, if we apply the additional assumption *N* → ∞ to (53) which already has *m* → 0, the pedigree effect becomes negligible.

#### Rare-migration limit

Here we consider *D* = 2 for fixed *N* and *m* ∈ (0, 1) as *α* → 0. We defer the detailed analysis to Supplementary Material Section 3. For this model, we write the transition matrix for the single-pair process as **Π**_*α*_ = **A** + *α***B**_*α*_, and the transition matrix for the two-pairs process as 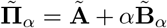, then apply Lemma 1 of Möhle (1998a) with time measured in units of 1*/α* generations.

Similar to the case of low migration, here two ancestral lines in initial state [••] will coalesce immediately regardless of the pedigree, because opportunities for migration are so infrequent in this limit. Under this rare-migration model, the conditional survival function of the pairwise coalescence time given an initial state [•][•] is *F*_*α*_(*t*) = ℙ(*τ* ^(*α*)^ *>* ⌊*t/α*⌋ | 𝒜 and our analysis gives

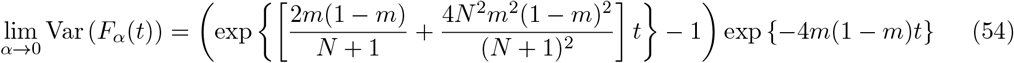

for *t* ≥ 0. Thus in the rare-migration limit as well, the coalescent process conditional on the population pedigree differs from the coalescent process marginal to the population pedigree.

In contrast to the low-migration regime of the previous subsection, if we consider the large population limit we obtain

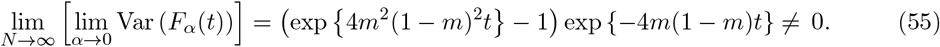

In this fourth case, the rare-migration limit, there is a persistent pedigree effect for initial state [•][•] even when we apply the additional assumption *N* → ∞.

### Robustness to diploid migration and dioecy

The four limit results in the previous subsections, Structured-coalescent limit, Many-demes limit, Low-migration limit and Rare-migration limit, were obtained under the simplifying assumptions of monoecy and gametic migration. In Supplementary Material Section 4 we apply the same four-fold analysis to a model of diploid migration. In Supplementary Material Section 5 we do the same for a model of dioecy with either gametic migration or diploid migration.

These extensions introduce additional states and transitions. Under diploid migration, for example, the two ancestral lines at a single locus can be in any of four possible states, not just the three considered so far: [•], [••] and [•][•]. Using parentheses to delineate individuals, we instead have [(•)] for one ancestral line in one individual in one deme, [(•)(•)] for two ancestral lines in two individuals in the same deme, [(•)][(•)] for two ancestral lines in two individuals in different demes, and [(••)] for two ancestral lines in the same individual in one deme. Further, the two ancestral lines in [(••)] must travel together. They cannot be split apart by migration.

Despite such important additional details, we show that our characterizations of the effects of the pedigree on pairwise coalescence times for each of the four possible limits also hold for diploid migration and dioecy. There are differences in how the various constants and rates which arise in the each of four limits depend on *N* and *m*. Interested readers are referred to Supplementary Material Sections 4 and 5. Overall, if the pedigree does (respectively, does not) retain its effect on the pairwise coalescence time in the limit under monoecy and gametic migration, then it also does (respectively, does not) under dioecy and diploid migration. This is summarized in Table 1, which also notes the cases in which the pedigree effect becomes negligible for larger deme sizes. Table S1 in Supplementary Material Section 6 displays the full expressions for the limiting variances of the conditional survival functions, denoted simply *V*_1_ through *V*_12_ in Table 1.

**Table 1:** Limiting variances of the conditional survival functions for the four limit models and two initial states of the sample. Of the twelve non-zero values, *V*_1_ to *V*_12_, demonstrating a pedigree effect for a particular case, eight tend to zero as *N* → ∞ (indicated in parentheses). For the cases involving diploid migration, [••] and [•][•] above are shorthand for [(••)] and [(•)][(•)]. The full expressions for *V*_1_ to *V*_12_ are given in Supplementary Material Section 6.

| Limit model | Initial state | Monoecy and gametic mig. | Monoecy and diploid mig. | Dioecy and gametic mig. | Dioecy and diploid mig. |
| --- | --- | --- | --- | --- | --- |
| Struct. coal. | $[\bullet\bullet]$ | 0 | 0 | 0 | 0 |
| | $[\bullet][\bullet]$ | 0 | 0 | 0 | 0 |
| Many demes | $[\bullet\bullet]$ | $V_1 (\rightarrow 0)$ | $V_2 (\rightarrow 0)$ | $V_3 (\rightarrow 0)$ | $V_4 (\rightarrow 0)$ |
| | $[\bullet][\bullet]$ | 0 | 0 | 0 | 0 |
| Low mig. | $[\bullet\bullet]$ | 0 | 0 | 0 | 0 |
| | $[\bullet][\bullet]$ | $V_5 (\rightarrow 0)$ | $V_6 (\rightarrow 0)$ | $V_7 (\rightarrow 0)$ | $V_8 (\rightarrow 0)$ |
| Rare mig. | $[\bullet\bullet]$ | 0 | 0 | 0 | 0 |
| | $[\bullet][\bullet]$ | $V_9$ | $V_{10}$ | $V_{11}$ | $V_{12}$ |

We anticipate that this robustness to different modes of reproduction and migration may hold more broadly, but we emphasize that all of the models we have considered are exceedingly simple. For example, in our model of monoecy and gametic migration all gametes are of the same type. In contrast, monoecious plants have “male” and “female” gametes with different migration patterns due to different dispersal mechanisms of pollen versus seeds (Levin and Kerster, 1974; Heuertz et al., 2003; Sork et al., 2015). In our model of dioecy, there are two mating types but what might make them different is not specified. In fact, no differences between them are posited in the model. McLaughlin et al. (2023) review recent work on sex differences and mating strategies. Miyagi et al. (2025) point out that the interpretation of patterns of genetic variation under sex-biased migration or admixture depends on assumptions about intrasexual variation.

### Simulation results and additional analysis

In this section, we illustrate variation among pedigrees using simulations of the pre-limiting model (in Structured population model with gametic migration) for the case of two demes, and present additional analysis of the many-demes model with large deme size. Simulations were implemented in the Julia programming language. Additional analysis was done with the aid of Mathematica (Wolfram Research, Inc., 2024). The associated code is included in [link TBD].

Figure 1 displays cumulative distribution functions, or CDFs, of coalescence times for the two initial states [••] and [•][•] on each of 1000 independently generated pedigrees under the pre-limiting model with *D* = 2 demes and allowing migration in every generation (*α* = 1). The CDF for a given sample on a given pedigree was estimated by simulating 10, 000 independent coalescence times, which are conditionally independent given the pedigree as would be true for 10, 000 unlinked loci in the sampled individual(s). The values of *N* and *m* were chosen so that *M* = 4*Nm* = 0.5 in order to illustrate convergence to the structured-coalescent limit. For the within-deme sample [••], Figure 1a shows that as *N* increases from 24 to 124 to 640, the variation in CDFs among pedigrees decreases, converging on the marginal limiting prediction from (16). Figure 1b displays the results for both the within-deme sample [••] and the between-deme sample [•][•] for the case *N* = 124, showing that both vary around their respective limiting prediction from (16).

**Figure 1:**
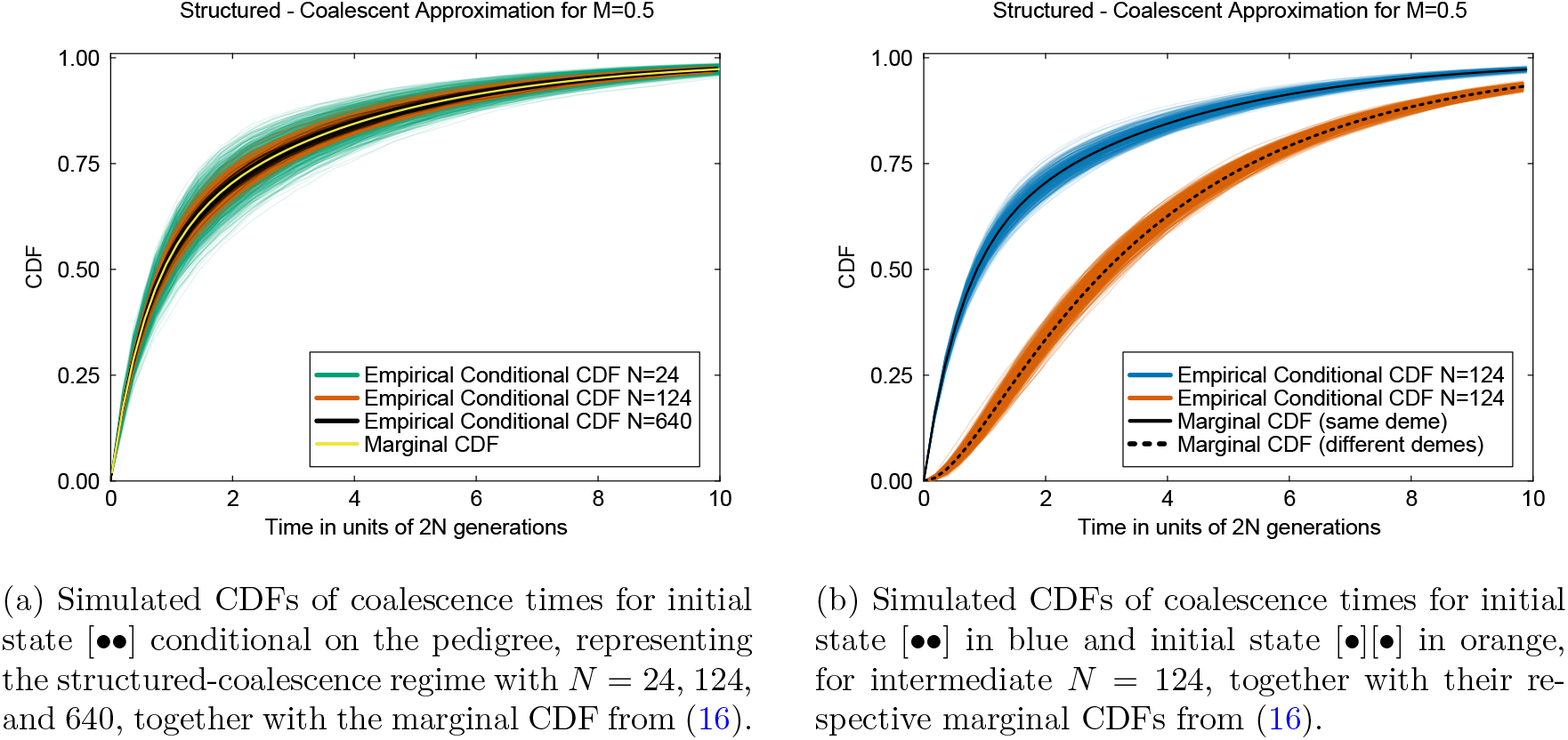
Simulations illustrating convergence to the structured-coalescence limit for *M* = 4*Nm* = 0.5.

Figure 2 shows the variability of CDFs of coalescence times for initial state [•][•] for two models with *D* = 2 and *N* = 24, but migration regimes representing the low-migration limit (Figure 2a) and the rare-migration limit (Figure 2b). In our pre-limiting model, *αm* is the average per-generation probability that a single genetic line migrates. This is held constant at *αm* = 0.0003 in both panels of Figure 2, but it is realized in two different ways: *α* = 1 and *m* = 0.0003 in the low-migration regime versus *α* = 0.001 and *m* = 0.3 in the rare-migration regime. As in Figure 1, CDFs were estimated by simulating 10, 000 conditionally independent coalescence times given each pedigree. Consistent with the variance among pedigrees tending to zero as *N* → ∞ in the low-migration limit but not the rare-migration limit, the CDFs in Figure 2b are more variable than those in Figure 2a. The CDFs in Figure 2b show dramatic jumps when migration events (of size *m* = 0.3, not shown) appear in the pedigree in specific past generations. For both regimes, the empirical averages of the 1000 CDFs are close to their limiting marginal predictions, from (S32) and (S52).

**Figure 2:**
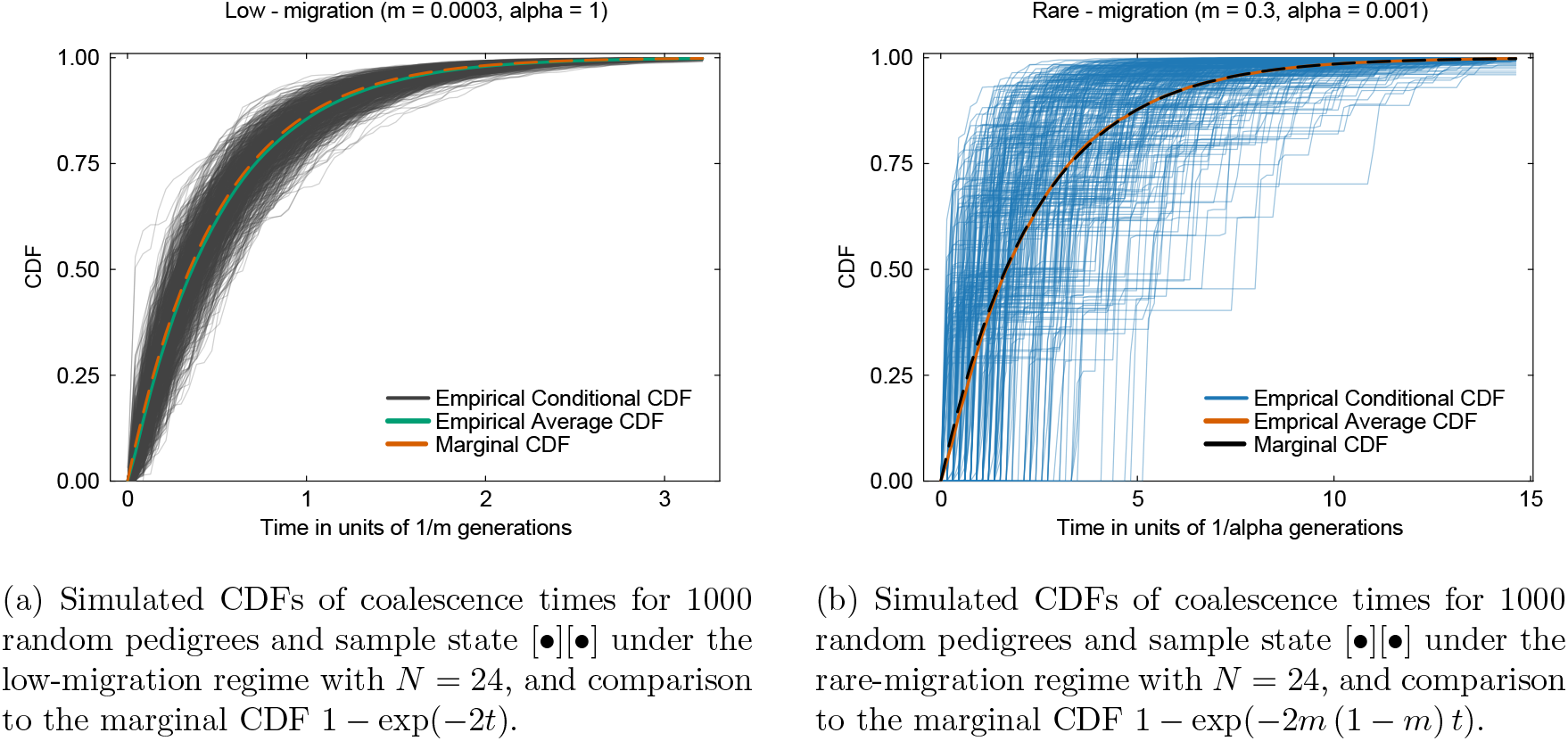
CDFs of coalescence times for two models with average migration probability *αm* = 0.003.

To illustrate the behavior of the many-demes limit, we do not display simulated CDFs of coalescence times but rather investigate the quantity 1 − 𝒰 − (1 − ℱ)^2^ in the expression (51) for the limiting variance under this model. As previously noted, this quantity and consequently the limiting variance tends to zero when we add the structured-coalescent assumption, *N* → ∞ for fixed *M* = 4*Nm*, to the many-demes limit. Here we consider the rate of approach of 1 − 𝒰 − (1 − ℱ)^2^ to zero when *N* is large. To leading order, this rate is equal to *g*(*M*)*/N* with

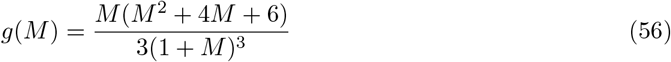

from the limit (S25). The function *g*(*M*) characterizes how quickly pedigree-induced fluctuations vanish as deme size increases, when *M* = 4*Nm* is held constant.

Figure 3 plots *g*(*M*) over a broad range of *M*. Fluctuations vanish fastest where *g*(*M*) has its maximum value, near *M* = 1. This is in line with intuition, that the structured-coalescent approximation will be most appropriate when *m* and 1*/N* are not too different in magnitude. For *M* close to zero (*m* ≪ 1*/N*), much larger values of *N* are required to make the pedigree effect disappear. In this case, the low-migration approximation may be a better choice than the structured-coalescent approximation. At the other extreme, for large *M, g*(*M*) approaches 1*/*3. Here, all effects of migration-mediated population structure become negligible, and we may anticipate connections to related results for well mixed populations. The quantity 1 − 𝒰 − (1 − ℱ)^2^ is the excess probability that neither of the two pairs of sampled gene copies at two unlinked loci coalesce in the recent ancestry described by **Ã** in (36), the excess being due to the shared pedigree. This same value of 1*/*3 was obtained as the leading-order coefficient of decay for the correlation of coalescence times for unlinked loci in a monoecious randomly mating population (Kogan et al., 2026, p. 50) as well as for the squared coefficient of variation of single-locus heterozygosity, also in a monoecious randomly mating population, which Weir et al. (1980) attributed to the pedigree.

**Figure 3:**
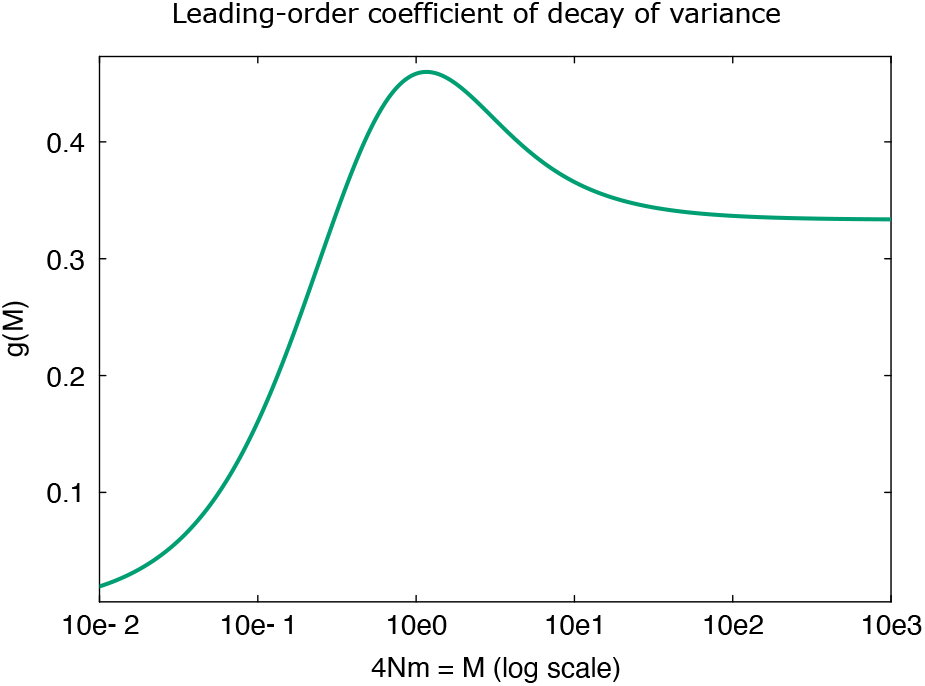
The rate of decay of the pedigree effect for large *N* and fixed *M* = 4*Nm* in the many-demes limit.

## Discussion

We have classified structured coalescent processes with migration in terms of how the shared population pedigree affects the distribution of pairwise coalescence times. We began with an idealized monoecious model of population subdivision and gametic migration, then extended our findings to diploid migration and dioecy. We quantified the effect of the pedigree in four different limits. Specifically, we asked whether the predicted distribution of pairwise coalescence times conditional on the pedigree differs from the one obtained by marginalizing or averaging over the pedigree.

Our idealized model has just four parameters: the number of demes *D*, the deme size *N*, the migration fraction *m*, and the migration probability *α*. The last of these, *α*, is the chance that any migration can occur at all, which, if it does, has migration fraction *m*. We did not find a pedigree effect in the structured-coalescent limit (*N* → ∞ with *m* proportional to 1*/N* for *D* = 2 and *α* = 1). But we did find a pedigree effect in the many-demes limit (*D* → ∞ for fixed *N, m* and *α* = 1), the low-migration limit (*m* → 0 for fixed *N, D* = 2 and *α* = 1) and rare-migration limit (*α* → 0 for fixed *N, m* and *D* = 2). These pedigree effects appear either for the within-deme sample [••] or the between-deme sample [•][•] depending on the particular limit as shown in Table 1, and they are similar in all the models. We also found that the pedigree effects in the many-demes limit and the low-migration limit are weak, in the sense that they become negligible as the deme size *N* increases. In the limits with a fixed number of demes, we have considered *D* = 2 for simplicity but the same conclusions are expected to hold for any *D* ≥ 2.

Regarding samples larger than two, we argue as follows. In cases where samples of size two show a pedigree effect, we may be confident larger samples will as well, because the properties of samples of size two are marginal properties of larger samples. It would be odd for them to disagree in this way. Indeed, Newman (2026) found a pedigree effect for larger samples in a generalized version of the rare-migration limit. In cases where samples of size two do not show a pedigree effect, additional work is needed to confirm that the full limiting conditional coalescent process for larger samples is identical to the traditional pedigree-averaged one. For a rigorous treatment of this in the structured-coalescent limit with an arbitrary number of demes, see Newman (2026). For the many-demes limit and the low-migration limit with the additional assumption *N* → ∞, we leave this as a conjecture, with some confidence from the results of Newman (2026) and analogous results for well-mixed populations (Tyukin, 2015; Alberti et al., 2025).

A finding of little or no pedigree effect on coalescence implies that inference methods developed for traditional coalescent models, which have all been derived by implicitly averaging over the pedigree, may be reinterpreted as properly accounting for the pedigree. This is useful because all loci in the genome share the same pedigree and traditional coalescent models are typically applied to data from multiple loci in a sample of individuals. A finding of an appreciable pedigree effect implies that new coalescent models and associated inference methods must be developed. Diamantidis et al. (2024) and Alberti et al. (2025) discuss some of the challenges involved.

Our results can be understood with reference to previous findings for populations without subdivision and migration, as well as to the general phenomenon of separation of time scales. In well-mixed outbred populations with biparental reproduction, the ancestral genetic lineages of any sample disperse to different individuals in *O*(1) generations (Möhle, 1998b; Birkner et al., 2013, 2018) and the pedigree ancestries of all individuals overlap completely in *O*(log_2_ *N*) generations (Chang, 1999). In contrast, the coalescent process takes *O*(*N*) generations.

Möhle (1998a,b) introduced the formal separation-of-time-scales approach to coalescent theory. The complementary formalism for the forward-time diffusion models of population genetics was introduced by Ethier and Nagylaki (1980). Möhle (1998a) treated self-fertilization, in which there is a non-negligible probability that two genetic lineages in a single individual coalesce due to self-fertilization in the recent ancestry of the individual. This is a property of sampled individuals, true even in the coalescent limit *N* → ∞, so self-fertilization is a fast-time-scale phenomenon. Möhle (1998b) treated dioecy, in which case the probability that two genetic lineages in a single individual coalesce in one generation is equal to zero, because the individual necessarily has two parents. This is also evident in the transition matrices (S120) and (S121) in Supplementary Material Section 5. Biparental reproduction with two mating types is another fast-time-scale phenomenon.

In the case of self-fertilization, in a single population of size *N*, the coalescent process marginal to the population pedigree differs from the Kingman coalescent process (Nordborg and Donnelly, 1997; Möhle, 1998a) and the coalescent process conditional on the population pedigree differs from the marginal one (Newman et al., 2025). Self-fertilization is an extreme form of inbreeding, which is known more generally to produce non-Kingman marginal coalescent processes (Severson et al., 2019, 2021) and may similarly always show pedigree effects in conditional coalescent models (Newman et al., 2025). None of the models we have considered here have any form of inbreeding, beyond the background chance 1*/N* of self-fertilization within demes in the monoecious case. So we do not consider inbreeding as an explanation of the pedigree effects we observe.

At the same time, our finding of a pedigree effect in the many-demes limit for a sample initially in state [••] but none for a sample initially in state [•][•] appears quite similar to the corresponding result for partial self-fertilization (Newman et al., 2025), only with demes of size *N* playing the role of diploid individuals (*N* = 2, in effect). Our ℱ in (29) is identical to *F*_*ST*_ in the island model of population structure (Wright, 1951). It quantifies the increase in relatedness due to subdivision, relative to that expected in a random-mating population of the same total size, and is generally considered a type of inbreeding coefficient. However, it is Wright’s *F*_*IS*_ that measures inbreeding owing to non-random mating strategies, such as partial self-fertilization, within local populations or demes. Our ℱ or *F*_*ST*_ can be made negligible simply by taking *N* → ∞, whereas *F*_*IS*_ is insensitive to population size. In all of our models, *F*_*IS*_ is equal to zero.

We consider finite population size or, more particularly finite deme size as the explanation of most of the pedigree effects on coalescence we observe. Such effects are expected in small populations even if the conditional and marginal coalescent processes are identical in the limit *N* → ∞ (Wakeley et al., 2012; Tyukin, 2015). An explanation in terms of finite *N* accords with our finding that the effects of the pedigree become negligible when we add the structured-coalescent assumption *N* → ∞ for fixed *M* = 4*Nm* to the many demes model and when we add the assumption *N* → ∞ to the low-migration model which already has *m* → 0.

With four parameters, *D, N, m* and *α*, there are a large number of possible limits we might have examined. The four we did examine were chosen because they are represented in the literature and they predict non-trivial levels of population structure, indicated by *F*_*ST*_ *>* 0. A fifth limit, which also appears in the literature but predicts *F*_*ST*_ = 0, is the strong-migration limit (Nagylaki, 1980; Notohara, 1993) here expressed most simply as *N* → ∞ with *m* ∈ (0, 1), *α* = 1 and arbitrary *D*. In the strong-migration limit, the movement of genetic lineages among demes is so frequent relative to their coalescence that the distribution of lineages among demes reaches stationarity, and the coalescent process reduces to a Kingman coalescent process with a rescaled rate or effective population size (Notohara, 1993).

With this in mind, we can check whether taking *N* → ∞ for *m* ∈ (0, 1) in the many-demes limit also makes the pedigree effect disappear for a sample initially in state [••]. From (49) and (51), this pedigree effect is revealed by the non-zero coefficient 1 − 𝒰 − (1 − ℱ)^2^ in the limiting variance of the conditional survival function, with ℱ as in (29) and 𝒰 obtained by solving (S7), (S8) and (S9) and setting 𝒰 = *u*_17_. Since the limiting conditional survival function becomes deterministic once in state [•][•], the variability comes only from the conditional probability it takes to reach state [•][•] from the initial state [••], whose expected value is 1 − ℱ and variance 1 − 𝒰 − (1 − ℱ)^2^. Letting *N* → ∞ makes this variance go to 0 as shown in Supplementary Material Section 1.2, and the effect of the pedigree does indeed become negligible. This holds not only with gametic migration in a monoecious population, but also with diploid migration and in a dioecious population (see Supplementary Material Sections 4, 5 and 6).

We conclude that the pedigree effect we observe in the many-demes limit is due to finite deme size, especially of the deme that initially contains the pair of ancestral lines. As such, we expect a persistent pedigree effect for other types of finely subdivided populations, for example metapopulations in which genetic exchange is mediated by extinction and recolonization rather than migration (Pannell and Charlesworth, 2000; Pannell, 2003). In fact, taking the limit *N* → ∞ may be seen as counter to the very idea of metapopulations, which are generally thought to be composed of small local subpopulations (Hanski and Gaggiotti, 2004).

Nordborg and Krone (2002) extended the strong-migration limit to allow migration fractions of order *O*(*N* ^−*θ*^) for some *θ* ∈ [0, 1), not just of order *O*(1), and called this fast migration. Hössjer (2011) adopted this terminology. Sagitov et al. (2010) studied coalescent processes in a related strong-migration limit for a haploid population model conditional on the migration fractions in each generation being determined by a random process, and found that it too reduced to a Kingman coalescent process. In light of these results, we might conjecture for example that a limit with *N* → ∞ for fixed *D, m* ∈ (0, 1) and *α* ∈ (0, 1) would yield the same coalescent process whether conditioning on or averaging over the pedigree, and that it would be a Kingman coalescent process.

The rare-migration limit is the only case in which we observe a pedigree effect on coalescence that we cannot attribute to finite deme size. In this limit, each migration event replaces a non-negligible fraction of a deme but the generations in which these events occur are separated by many generations without gene flow, with complete isolation between demes. Similarly, the times and sizes of large reproduction events have a strong effect on coalescence times in a single well mixed population even in the limit *N* → ∞ (Diamantidis et al., 2024; Alberti et al., 2025). Taking these two results together suggests that pedigree effects on coalescence will be observed whenever ‘large’ events happened with some frequency on the coalescent time scale. Here a ‘large’ event is one which affects multiple ancestral genetic lineages or multiple pairs of ancestral genetic lineages in the same deme or in different demes. This is precisely what is shown in Newman (2026).

Such large, rare migration events can also be referred to as pulse admixture or introgression (Iasi et al., 2021; Thawornwattana et al., 2023). Our findings, together with those of Newman (2026), of a pedigree effect in this case means that coalescence depends on the particular times and sizes of these events. In fact, pulse admixture and introgression are typically treated as having happened at fixed times in the past, even in situations where they are referred to as recurrent (Buzbas and Verdu, 2018; Ferreira et al., 2020; Di Santo et al., 2023). Our results and those of Newman (2026) also support the use of prior models for the times and sizes of such events in Bayesian approaches to inference, as discussed in Diamantidis et al. (2024) for the case of large reproduction events.

In contrast to the rare-migration limit, we note that the reason the pedigree effect becomes negligible as *N* → ∞ in the low-migration limit is that migration events will almost surely involve single individuals. Although pedigrees will contain many generations without any migration events, when *N* is large it will be unlikely for genetic lineages to trace back to those single migrant individuals in generations where migration has occurred. Conditionally independent genetic lineages will randomly use different migration events, making the distribution of pairwise coalescence times among unlinked loci insensitive to the details of the pedigree.

## Supporting information

associated_code

## Acknowledgements

We thank Yichen Si for helpful calculations and discussions in the initial stages of this project. Terry Easlick and Sabin Lessard were supported in part by Natural Sciences and Engineering Research Council of Canada (NSERC) Discovery Grant 8833 to Sabin Lessard.

## Supplementary Material

Lessard S, Easlick T, Wakeley J (2026) A classification of structured coalescent processes with migration, conditional on the population pedigree. …

### 1 Many-demes limit, additional calculations

#### 1.1 Fast-process absorption probabilities of the many-demes limit

As in the main text, let *u*_*i*_ denote the absorption probability in the coalescent state 1 given initial state *i*. From (41), we have *u*_1_ = 1, *u*_3_ = *u*_4_ = ℱ, and *u*_2_ = *u*_5_ = *u*_6_ = *u*_7_ = *u*_8_ = *u*_9_ = 0. Using these and the block structure of **Ã** in (37), we can solve for the remaining *u*_*i*_ with *i* ∈ {10, …, 17} using first-step analysis.

There are 35 non-zeros entries in **Ã**_21_ and 26 non-zeros entries in **Ã**_22_. We do not detail these here but they are straightforward to compute. In short: each gamete is a migrant with probability *m* independently of all other gametes; each gamete traces back to one of the *N* parental individuals in its deme uniformly and independently of all other gametes; then the gene copy or copies in each gamete trace back to the gene copies in the parent according to Mendel’s laws.

For example, consider state 12, or [••][○○]. In the process described by **Ã**, which recall has transition probabilities of order *O*(1) in the limit *D* → ∞, state 12 cannot be reached from any other state. But states 1, 3 and 4 can be reached in one step from state 12. The transition probability from state 12 to state 1 can be written

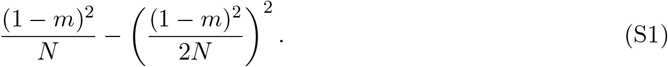

This can be derived by accounting for all possibilities using the rules in the previous paragraph, for example there being zero, one, two, three or four migration events. Alternatively, it can be obtained by noting that the pairs [••] and [○○] in state 12 independently coalesce with probability (1 − *m*)^2^*/*(2*N*), and the transition to state 1 entails the coalescence of at least one pair.

Considering all possible single-step outcomes for state 12, we have

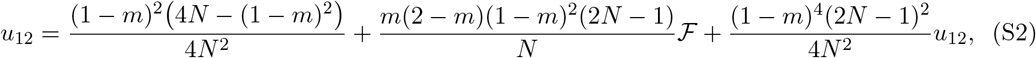

in which the first term on the right-hand side is another way of writing (S1) and the second term is the one-step transition probability to either state 3 or state 4 times the probability of absorption in state 1 from those states. Solving (S2) gives *u*_12_ = 1 − (1 − ℱ)^2^.

In the same way, we have

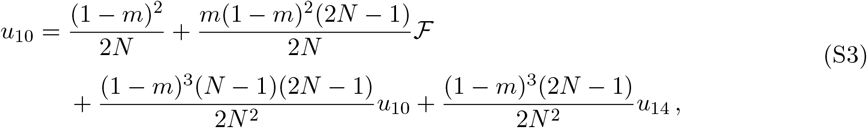

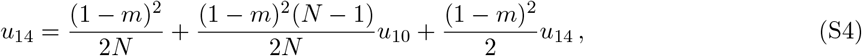

which give *u*_10_ = *u*_14_ = ℱ, and

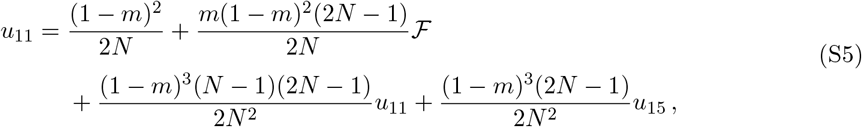

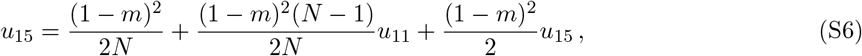

which give *u*_11_ = *u*_15_ = ℱ.

The absorption probabilities *u*_13_, *u*_16_ and *u*_17_ for [• • ○○], 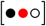, and 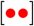 are considerably more complicated. Using *u*_1_ = 1, *u*_3_ = *u*_4_ = *u*_10_ = *u*_11_ = *u*_14_ = *u*_15_ = ℱ, and *u*_2_ = *u*_5_ = *u*_6_ = *u*_7_ = *u*_8_ = *u*_9_ = 0, we have

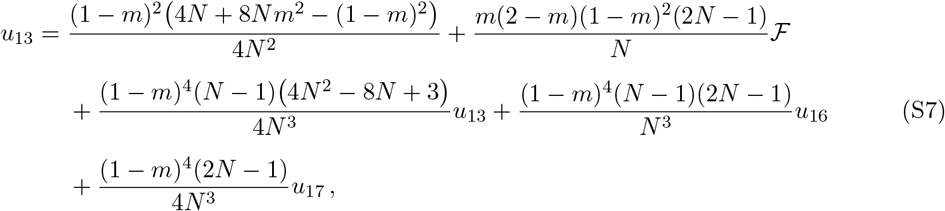

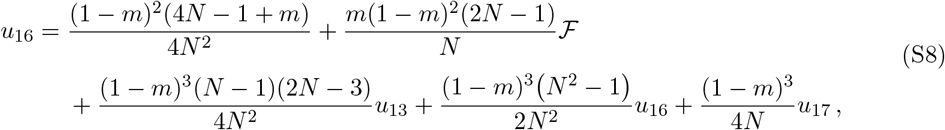

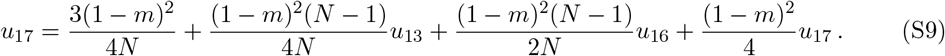

The first term on the right-hand sides of (S7), (S8) and (S9) are the single-step transition probabilities to state 1 from states 13, 16 and 17, respectively. The terms multiplied by ℱ in (S7) and (S8) are the total single-step probabilities of going from states 13 and 16 to any of the states 3-4, 10-11, or 14-15, from which the probability of eventual absorption in state 1 is equal to ℱ. The remaining terms correspond to transitions among states 13, 16 and 17. Solving (S7), (S8) and (S9) for *u*_13_, *u*_16_, *u*_17_ results in the unwieldy expressions shown in Supplementary Material Section 1.1. Using these unwieldy expressions, confirmation that the quantity 1 − *u*_17_ − (1 − ℱ)^2^ is indeed not equal to zero for any finite *N* is given in Supplementary Material Section 1.3.

The Mathematica notebook `q17manydemes.nb` included in `associated_code.zip` gives the expression for the fixation probabilities *u*_13_, *u*_16_, *u*_17_ on the fast time scale in the many-demes limit of a monoecious population with gametic migration, and further displays the structured-coalescent and strong-migration limits of *u*_17_, which for convenience we also call 𝒰, in the main text.

#### 1.2 Adding the structured-coalescent assumption to the many-demes limit

Here we consider *m* = *M/*(4*N*) as *N* → ∞ with *M >* 0 kept constant, using Lemma 1 of Möhle (1998a). The transition matrix in the limit of an infinite number of demes, that is **Ã** in (36) and (37), can be expressed as

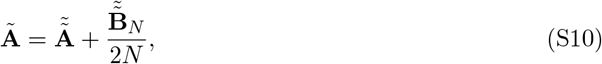

where

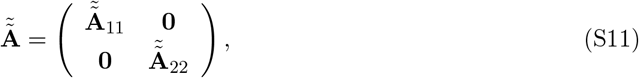

is the transition matrix when *N* = ∞ and *m* = 0 with

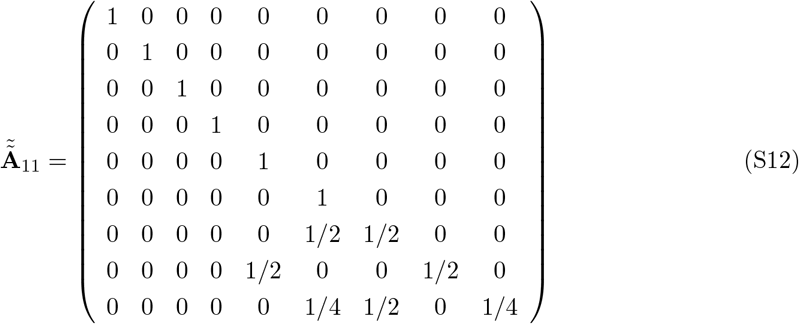

and

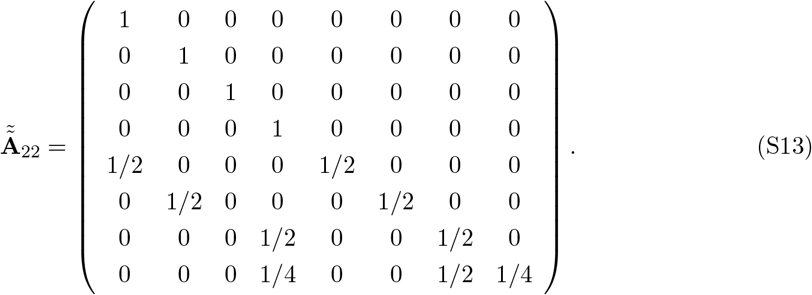

Moreover, we have

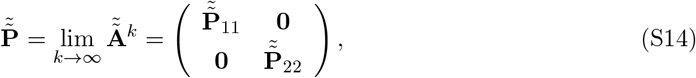

where

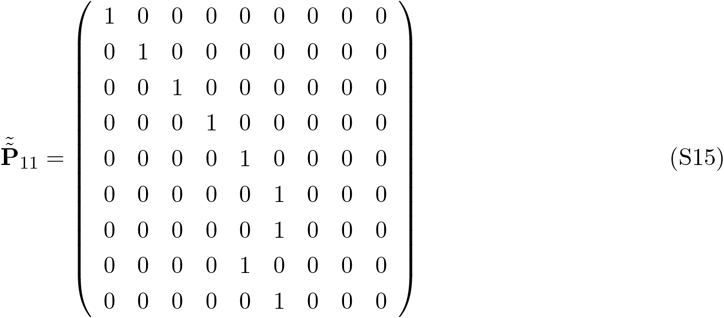

and

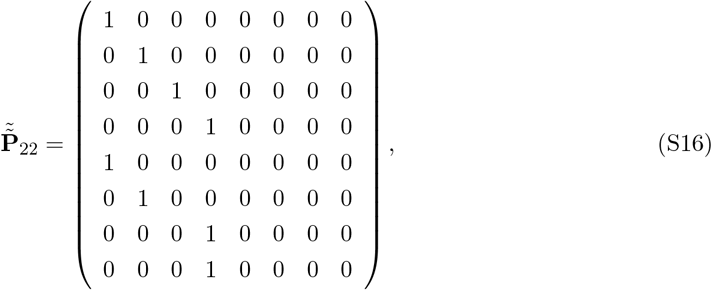

while the rows 10, 11, 12, 13 of

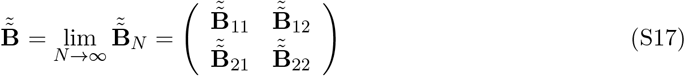

are given by

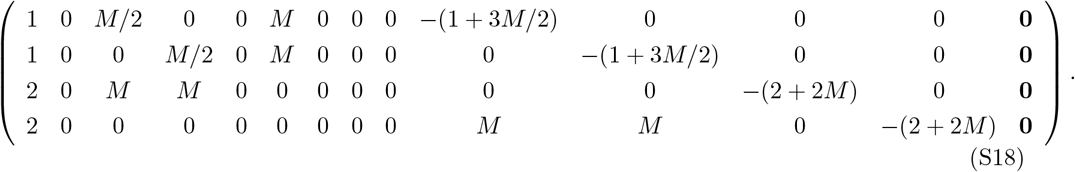

These are also the rows 10, 11, 12, 13 of

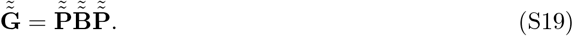

Applying Lemma 1 of Möhle (1998a), we have

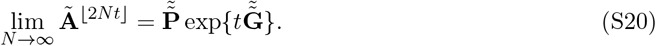

In the limit as *N* → ∞ with 2*N* time steps as unit of time, there are instantaneous transitions from states in {14, 15, 16, 17} to states in {10, 11, 12, 13} from which the rates of change are given in (S18).

Conditioning on the first change of state, the probabilities of fixation into state 1 starting from states in {10, 11, 12, 13} in the limit as *N* → ∞ are given by

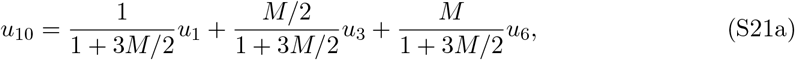

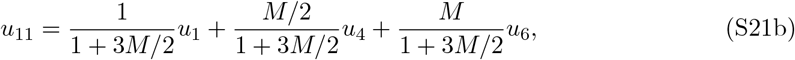

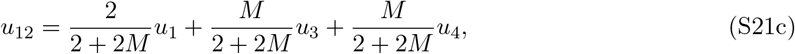

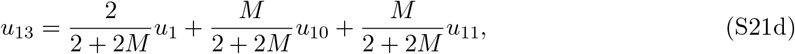

where *u*_1_ = 1, *u*_3_ = *u*_4_ = 1*/*(1 + *M*) and *u*_6_ = 0. We obtain

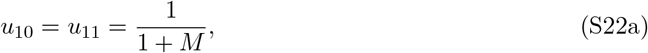

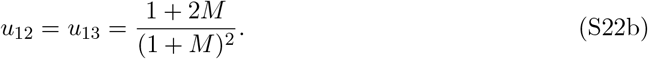

We conclude that

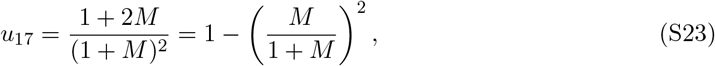

where *M/*(1 + *M*) = lim_*N*→∞_(1 − ℱ) with ℱ given in (29).

#### 1.3 For all finite N, 1 − u_17_ − (1 − ℱ)^2^ is not equal to zero

We employ Bernstein polynomial representations to further analyze 1 − *u*_17_ − (1 − ℱ)^2^. For a fixed degree *d*, the Bernstein polynomial basis on [0, 1] is given by

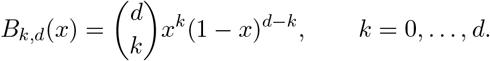

These basis polynomials span the space of polynomials of degree at most *d*, and thus any polynomial *p*(*x*) with deg(*p*) ≤ *d* admits the exact representation

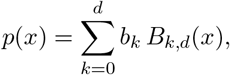

for suitable coefficients *b*_*k*_. Although the quantity 1 − *u*_17_ − (1 − ℱ)^2^ depends on *N* and *m*, we hereafter apply the Bernstein representation in the variable 0 *< m <* 1, treating *N* ≥ 4 as a parameter. The Mathematica notebook provides rational expressions made of polynomials in *m* for each fixed *N*, and hence we may write

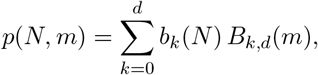

where 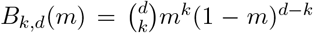 and the coefficients *b*_*k*_(*N*) depend polynomially on *N*. The above machinery allows us to write

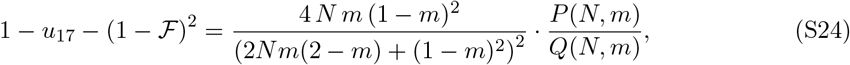

where

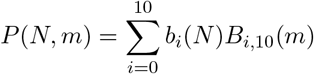

is a Bernstein polynomial with coefficients

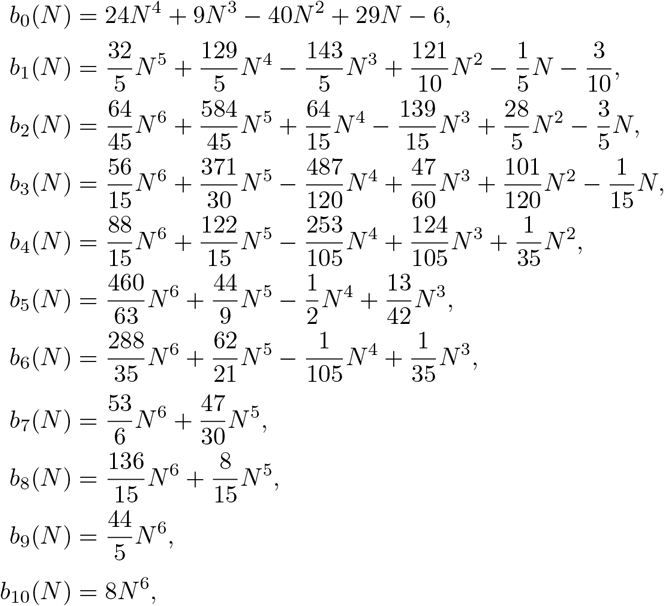

and 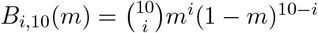. Similarly,

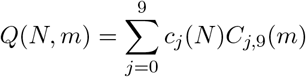

is a Bernstein polynomial with coefficients

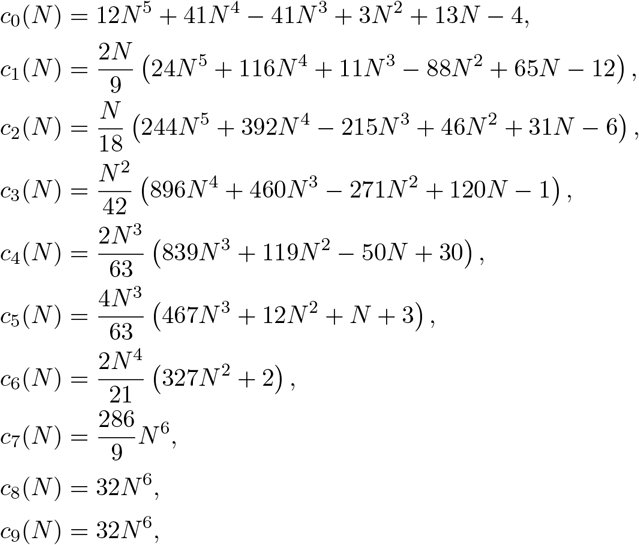

and 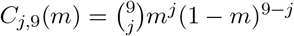.

First note that all of the above Bernstein basis polynomials are positive on (0, 1). Secondly, a direct check shows that *b*_*i*_(4) *>* 0 for *i* = 0, …, 10 and *c*_*j*_(4) *>* 0 for *j* = 0, …, 9. Each *b*_*i*_(*N*) and *c*_*j*_(*N*) is a polynomial in *N* with positive leading coefficient. Momentarily treating as real-valued polynomials for *x* ∈ ℝ, a quick calculation shows that 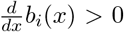 and 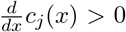 for all *x* ∈ [4, ∞). Hence *b*_*i*_(*x*) *>* 0 and *c*_*j*_(*x*) *>* 0 for all *x* ≥ 4, and in particular *b*_*i*_(*N*) *>* 0 and *c*_*j*_(*N*) *>* 0 for all *N* ≥ 4. Thus it follows that *P* (*N, m*) *>* 0 and *Q*(*N, m*) *>* 0 for all *N* ≥ 4 and 0 *< m <* 1. Additionally, it is not hard to verify that

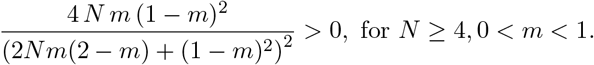

Therefore, the quantity 1 − *u*_17_ − (1 − ℱ)^2^ is not equal to zero. Moreover, these results concretely illustrate that infinite global population yet finite deme size is not sufficient to suppress pedigree effects; namely, such effects disappear only when deme sizes also grow without bound.

#### 1.4 Limiting behaviour of (1 − u_17_ − (1 − ℱ)^2^) as N → ∞ with 4Nm = M

We analyze the limiting behaviour of the many-demes model under the structured-coalescent assumption. In this regime, pedigree effects vanish at a rate inversely proportional to deme size. Using the results of the previous subsection, and taking the limit *N* → ∞ with 4*Nm* = *M* held fixed, we obtain

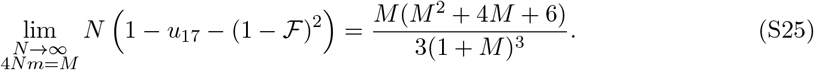

A derivation of (S25) using the Bernstein polynomials with coefficients given above is provided in the Mathematica notebook `manydemes_variance_limit_behaviour.nb` included in `associated_code.zip`.

### 2 Low-migration limit analysis

In this section, we will consider *D* = 2 demes of fixed size *N* and let *m* → 0 for *α* = 1. The possible states for the ancestral lines of a pair of genes at a single locus in an individual are the same as in the main text: [•] for only one in one deme, [••] for two in the same deme, and [•][•] for two in different demes. As for the transition matrix from one generation to the previous one, it can be expressed as

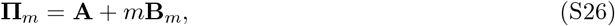

where

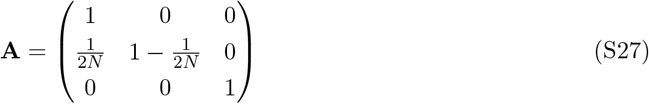

and

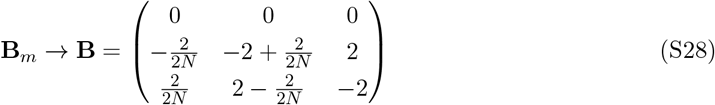

as *m* → 0. By Lemma 1 of Möhle (1998a), we have

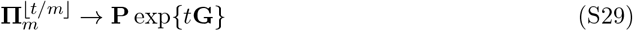

as *m* → 0, where

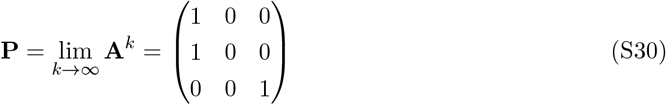

and

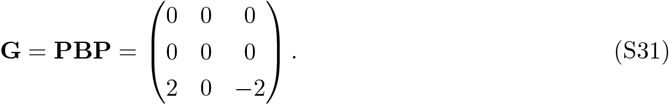

Equation (S30) shows that ancestral lines that start in initial state [••] coalesce instantaneously on the new time scale as *m* → 0 regardless of the details of the pedigree. For ancestral lines that start in initial state [•][•], the conditional survival function *F*_*m*_(*t*) = ℙ(*τ* ^(*m*)^ *>* ⌊*t/m*⌋ | 𝒜 satisfies

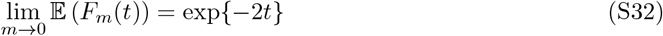

for *t* ≥ 0.

Now, considering the ancestral lines that start from two pairs of genes at unlinked loci labeled 1 and 2 in an individual, the possible states for the ancestral sequences are the same as in the previous subsection but they are ordered differently as below:

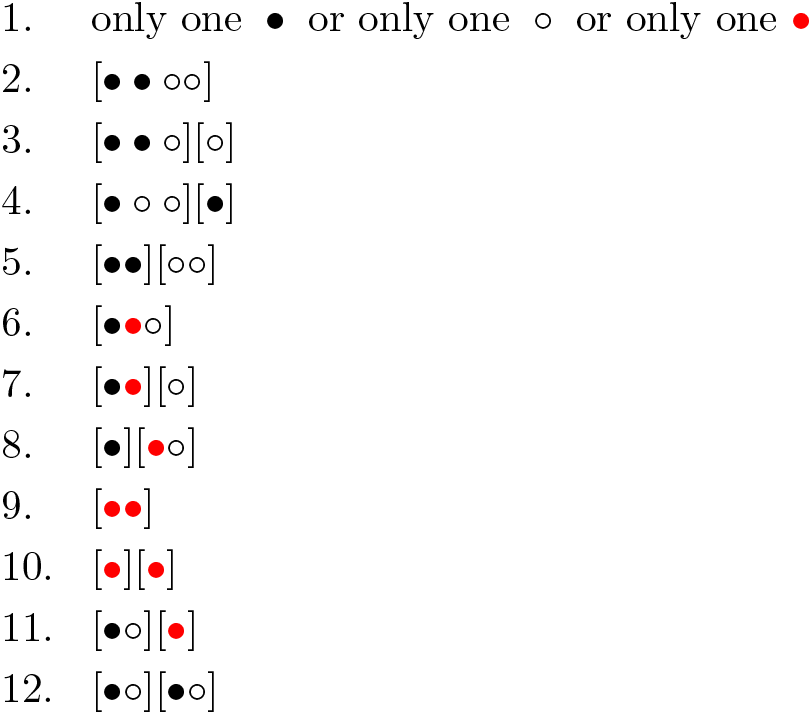

With respect to these states, the transition matrix from one generation to the previous one takes the form

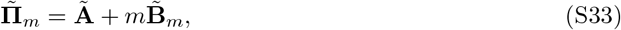

where **Ã** is the transition matrix in the absence of migration. In this case, there will be absorption into state class 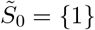 from any initial state in 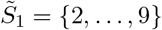 as a result of coalescence events. On the other hand, from any initial state in 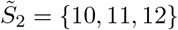 the ancestral lines in each deme will follow a Markov chain on the state space 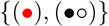, independently of the ancestral lines in the other deme, whose probabilities of transition by recombination or coalescence are given by

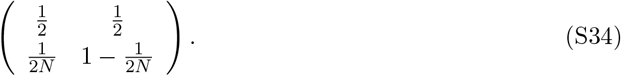

The stationary distribution of this chain is (1*/*(*N* + 1), *N/*(*N* + 1)). With respect to the states in 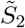 the entries of **Ã** are given by

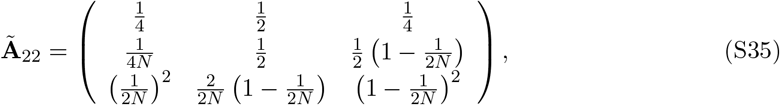

which satisfies

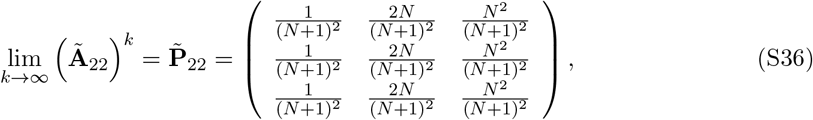

owing to the ergodic theorem for discrete-time Markov chains on a finite state space (see, e.g., Karlin and Taylor (1975)). Moreover, with respect to the state sets 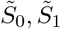 and 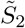, we have

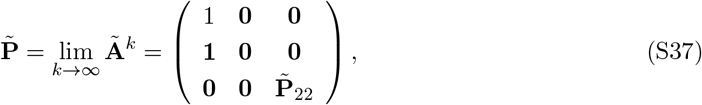

where **0** and **1** are matrices of zeros and ones, respectively. With respect to the same state sets, the rate matrix for changes that involve migration events in the limit as *m* → 0 takes the form

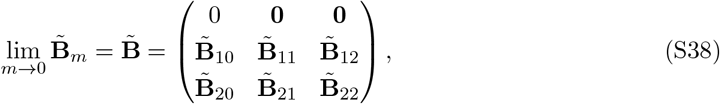

where

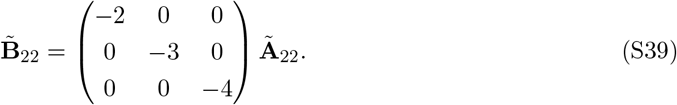

Lemma 1 of Möhle (1998a) guarantees that

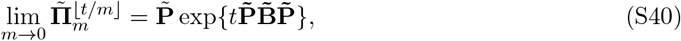

where

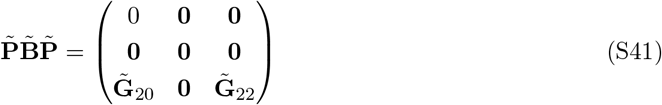

with

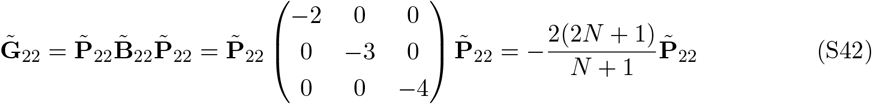

and

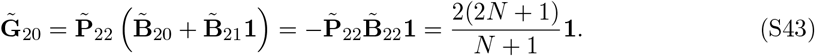

This uses the fact that the entries on each row of 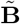. We conclude that

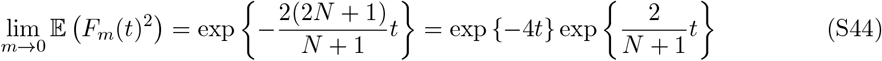

for *t* ≥ 0. Combining this with our previous result yields

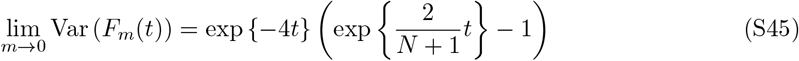

for *t* ≥ 0, which we note tends to zero as *N* → ∞.

### 3 Rare-migration limit analysis

In this section, we will consider *D* = 2 for fixed *N* and *m* ∈ (0, 1) as *α* → 0. The possible states for the ancestral lines of a pair of genes at a single locus in an individual are the same as the prior section: [•] for only one in one deme, [••] for two in the same deme, and [•][•] for two in different demes. In similar fashion as before, we can express this transition matrix as

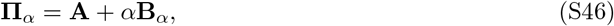

where

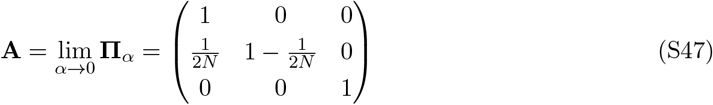

and

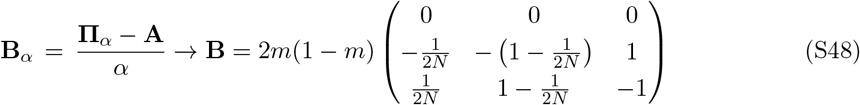

as *α* → 0. We again utilise Lemma 1 of Möhle (1998a) and obtain

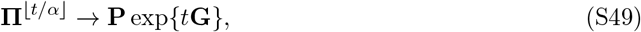

as *α* → 0, where

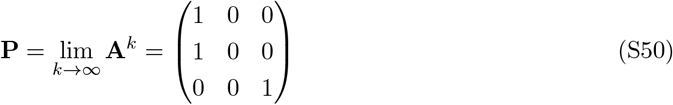

and

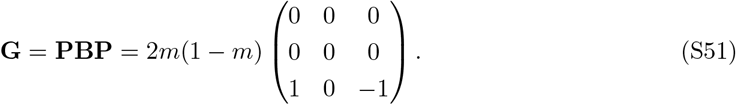

Equation (S50) implies that ancestral lines starting in initial state [••] coalesce instantaneously on the new time scale in the limit as *α* → 0. For ancestral lines that start from initial state [•][•], the survival function *F*_*α*_(*t*) = ℙ(*τ* ^(*α*)^ *>* ⌊*t/α*⌋ | 𝒜 satisfies

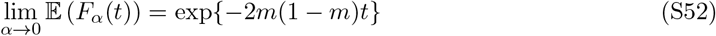

for *t* ≥ 0.

We again consider the ancestral lines that start from two pairs of genes at unlinked loci labeled 1 and 2 in an individual, and the same ordering of the possible states for the ancestral sequences as in the previous subsection:

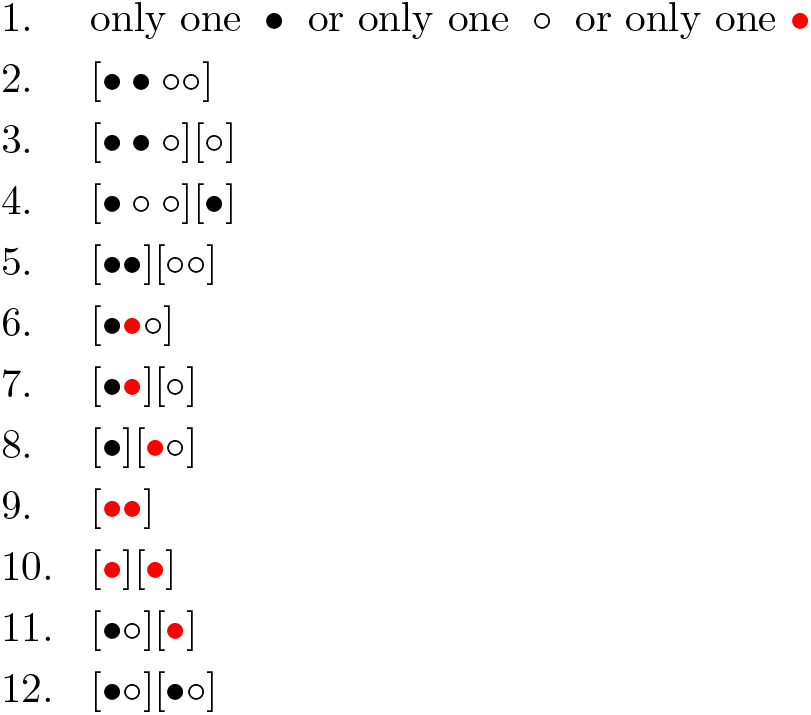

With respect to these states, the transition matrix from one generation to the previous one takes the form

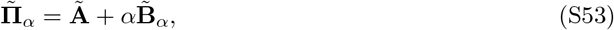

where **Ã** is again the no-migration transition matrix from the low-migration regime. States in 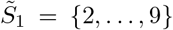 absorb into 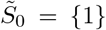 via coalescence, while in 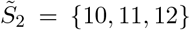 each deme’s ancestral lines evolve independently on 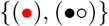 with recombination and coalescence transition probabilities given by

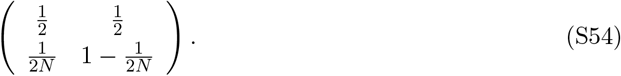

With respect to these subsets of states, we have as previously

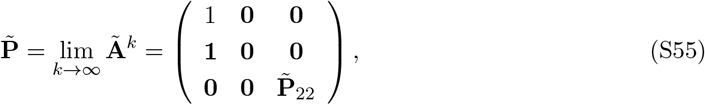

where

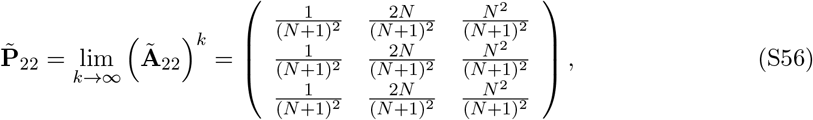

while **0** and **1** are matrices of zeros and ones, respectively. With respect to the same state sets, the rate matrix for changes that involve migration events in the limit as *α* → 0 takes the form

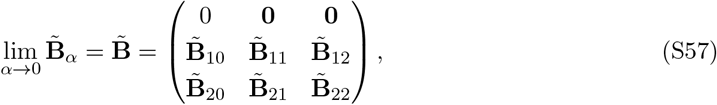

where

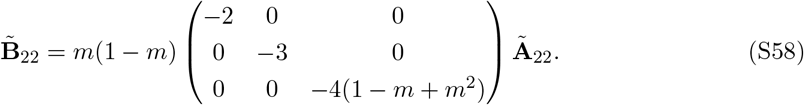

Notice that 2*m*(1 − *m*) = 1 − (1 − *m*)^2^ − *m*^2^, while we have 3*m*(1 − *m*) = 1 − (1 − *m*)^3^ − *m*^3^ and 4*m*(1 − *m*)(1 − *m* + *m*^2^) = 1 − (1 − *m*)^4^ − *m*^4^ − 2*m*^2^(1 − *m*)^2^.

Lemma 1 of Möhle (1998a) guarantees that

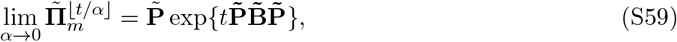

where

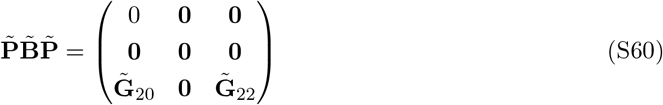

with

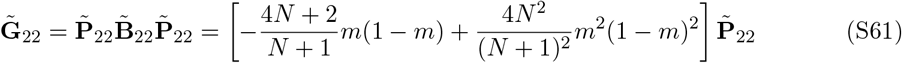

and

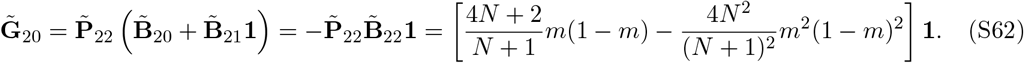

Therefore, we obtain

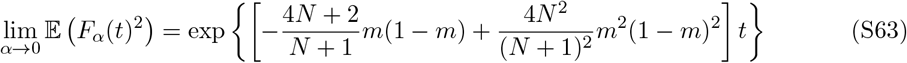

for *t* ≥ 0. Moreover, combined with the earlier results, Var (*F*_*α*_(*t*)) tends to

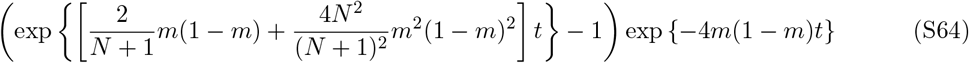

as *α* → 0 for *t* ≥ 0.

### 4 Monoecious population with diploid migration

In this section, we consider a monoecious diploid population as in the main text but with migration of offspring instead of migration of gametes. Under this assumption, we have to distinguish between ancestral lines in the same individual and ancestral lines in different individuals in the same deme.

The possible states for two ancestral lines at a single locus are: [(•)] for one in one individual in one deme, [(•)(•)] for two in two individuals in the same deme, [(•)][(•)] for two in two individuals in different demes, and the additional state [(••)] for two in the same individual in one deme. The transition matrix with these states in this order is given by

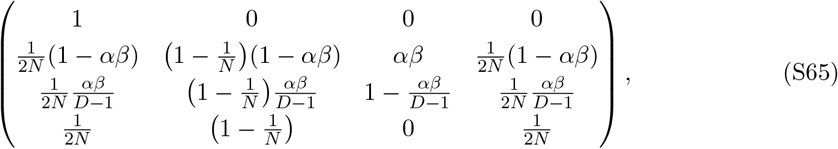

in which

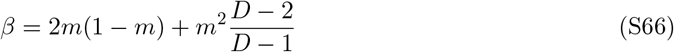

represents the conditional probability for two offspring chosen at random in the same deme to come from two different demes given that migration occurs, and *β/*(*D* − 1) the conditional probability for two offspring in different demes to come from the same deme.

#### 4.1 Structured-coalescent limit

In this subsection, we consider *D* = 2 demes of large size *N* and a fixed deme-scaled migration fraction *m* = *M/*(4*N*) with probability *α* = 1 in each generation. The transition matrix in (S65) for a pair of ancestral lines at a single locus can be written in the form

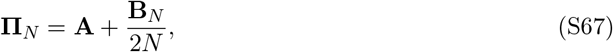

where

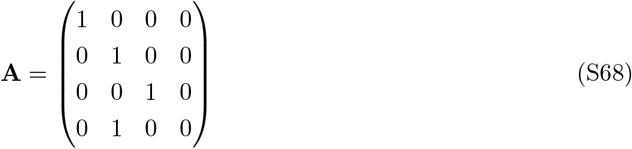

and

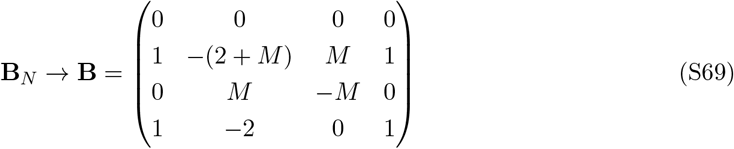

as *N* → ∞. Using Lemma 1 of Möhle (1998a), we have

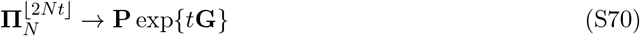

as *N* → ∞, where

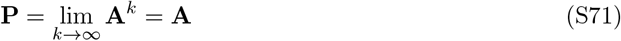

and

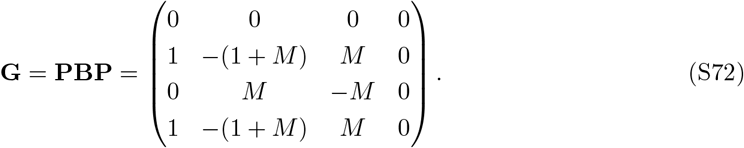

Therefore, in the limit *N* → ∞ and starting from state [(••)], there is an instantaneous transition to state [(•)(•)] and this is followed by a continuous time Markov chain on the states [(•)], [(•)(•)] and [(•)][(•)] with rate matrix given in (9), i.e., just as in the case of gametic migration.

For two pairs of ancestral lines at two unlinked loci, the state space *S* can be decomposed into the following subsets: *S*_1_ = {1, …, 6} as in Structured-coalescent limit for fewer than four ancestral lines or four ancestral lines in four different individuals, *S*_2_ for four ancestral lines in the same deme except state 2, *S*_3_ for two ancestral lines at locus 1 and one at locus 2 in one deme and another ancestral line at locus 2 in a different deme except state 3, *S*_4_ for two ancestral lines at locus 2 and one at locus 1 in one deme and another ancestral line at locus 1 in a different deme except state 4, *S*_5_ for one ancestral line at locus 1 and one at locus 2 in each of two different demes except state 5, and *S*_6_ for two ancestral lines at locus 1 in one deme and two ancestral lines at locus 2 in a different deme except state 6. The transition matrix with respect to these subsets can be written

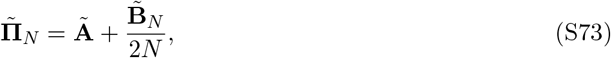

where

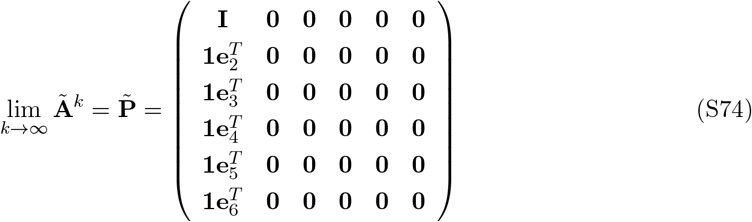

with **1** representing a column vector of all ones and 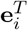 being the *i*-th row vector of the 6-dimensional identity matrix **I** for *i* = 1, …, 6. Further, the first 6 rows of 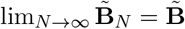 are given by

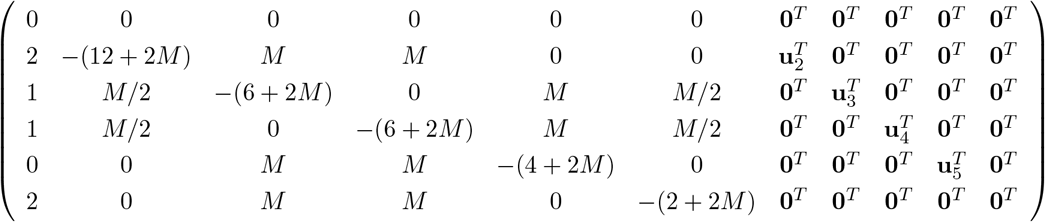

with 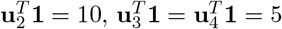 and 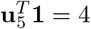. In this case, Lemma 1 of Möhle (1998a) yields

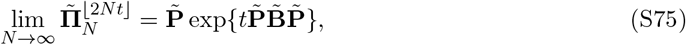

where the entries of 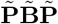 corresponding to the states in *S*_1_ are given by

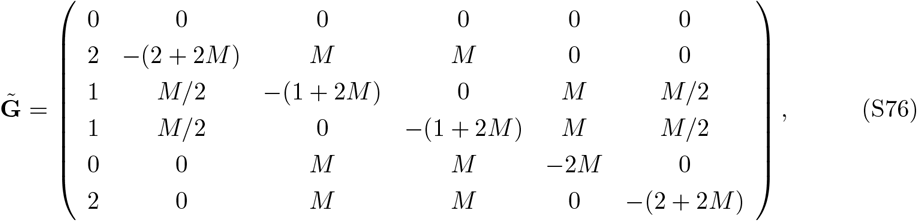

which is the same as the rate matrix in (22) for the case of gametic migration.

We can conclude that the results in Structured-coalescent limit apply *mutadis mutandis*.

#### 4.2 Many-demes limit

For *N* and *m* fixed with *α* = 1 as *D* → ∞, the transition matrix for a pair of ancestral lines at the same locus denoted by **Π**_*D*_ satisfies

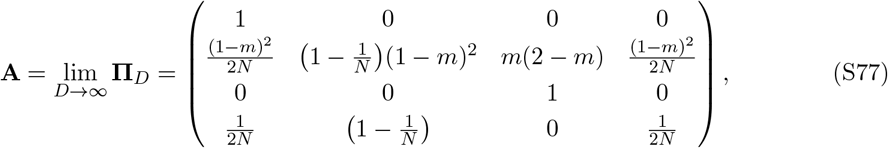

from which

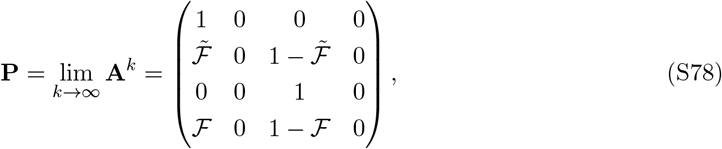

where

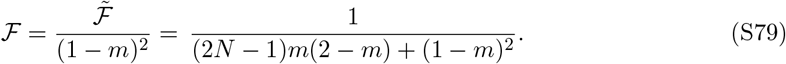

Furthermore,

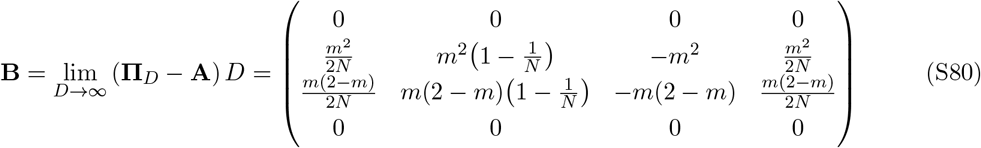

and

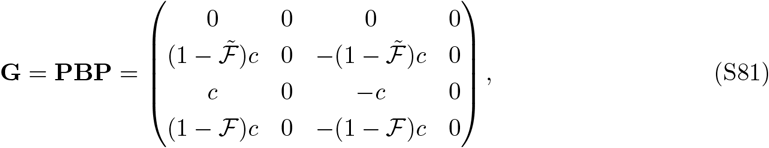

where

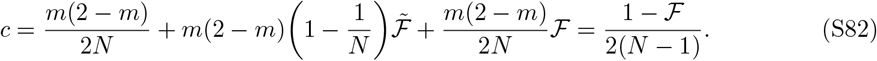

This is the rate of coalescence of two ancestral lines starting in individuals in different demes with *D* generations as unit of time as *D* → ∞. This differs slightly from the corresponding result (1 − ℱ)*/*(2*N*) in (32) for the case of gametic migration. Starting from state [(••)] and owing to Lemma 1 of Möhle (1998a), which ensures that lim_*D*→∞_ **Π**_*D*_^⌊*tD*⌋^ = **P** exp{*t***G**}, we have

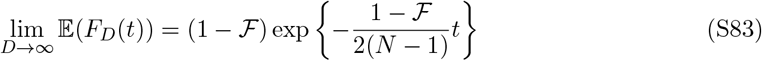

for *t* ≥ 0, which differs from the corresponding result (35) for the case of gametic migration only by the replacement of 2*N* with 2(*N* − 1) in the rate of coalescence for *t >* 0.

As for two pairs of ancestral lines at two unlinked loci, the transition matrix from one generation to the previous one can be expressed in the form

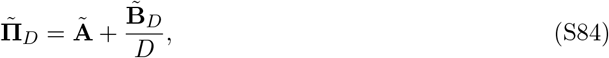

where **Ã** is the transition matrix in the limit of an infinite number of demes. In this limit, all states but two are transient. The state with only one ancestral line at any one of the loci and the state with the four ancestral lines in all different individuals in all different demes are absorbing. Call these states 1 and 2, respectively. Taking *D* generations as unit of time and letting *D* go to infinity, there are instantaneous transitions to these states 1 and 2 from any other state. Moreover, once in state 2, the rate of transition to state 1 is the rate at which two individuals carrying two ancestral lines at the same locus come from the same deme, times the probability that the two ancestral lines come from the same gene in the same parent, plus the probabilities that they come either from the two genes in the same parent or from two genes in different parents, and coalesce before migrating in the limiting process. This gives the rate

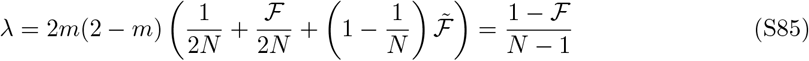

which is twice the rate in (S82) and which we note differs slightly from the corresponding result for gametic migration, (1 − ℱ)*/N* in (48). With diploid migration, therefore, starting with genes at two unlinked loci on two chromosomes in different demes, represented by 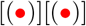, there is an instantaneous transition to state 2 and this is followed by a transition to the fixation state 1 at rate (1 − ℱ)*/*(*N* − 1).

Starting with initial state 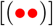, so that both pairs of lines start from the same chromosomes in the same deme, however, there will be an instantaneous transition to either the fixation state 1 with some probability 𝒰, or to state 2 with the complementary probability 1 − 𝒰 from which the transition to the fixation state 1 will occur at rate (1 − ℱ)*/*(*N* − 1). This 𝒰 corresponds to *u*_17_ in Many-demes limit and in Section 1.1. We do not attempt to derive it here but we anticipate that like *u*_17_ it will not in general be equal to 1 − (1 − ℱ)^2^, which cf. (51) would be the value that would make lim_*D*→∞_ Var(*F*_*D*_(*t*)) here equal to zero.

However, in the limit of a large deme size with *M* = 4*Nm* kept constant, ancestral lines cannot stay in the same individual with probability one and we have

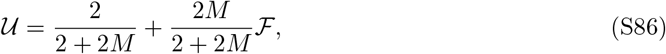

where here we use

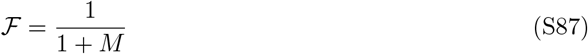

to represent the limiting *N* → ∞ value of (S79) with *M* = 4*Nm* kept constant. This yields

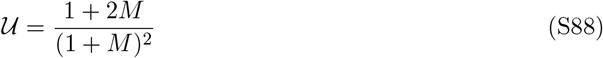

which is identical to the corresponding value for gametic migration (S23) derived in Section 1.1. We conclude similarly that, under diploid migration, the effect of conditioning on the pedigree becomes negligible with the addition of the structured-coalescent assumption to the many-demes model.

#### 4.3 Low-migration limit

For *D* = 2 demes and a small migration fraction *m* with probability *α* = 1 in each generation, the transition matrix in (S65) for a pair of ancestral lines at a single locus can be written in the form

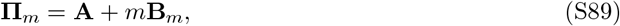

where

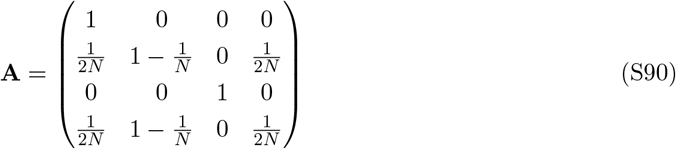

and

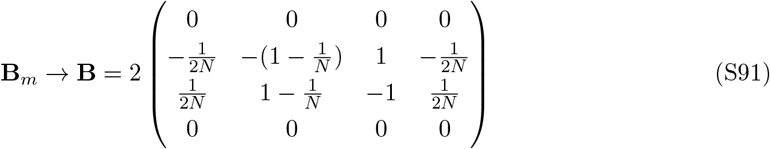

as *m* → 0. By Lemma 1 of Möhle (1998a), we have

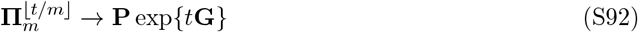

as *m* → 0, where

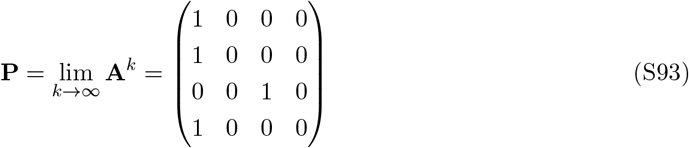

and

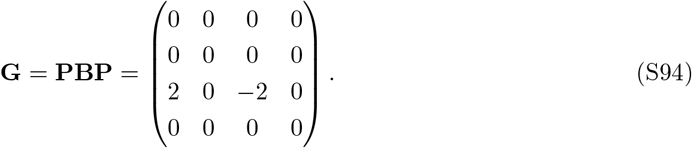

Therefore, starting from the state [(•)][(•)], the survival function satisfies

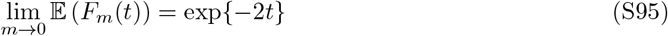

for *t* ≥ 0, which is identical to the corresponding result (S32) for gametic migration. On the other hand, starting from the initial state [(••)], there is instantaneous coalescence in the limit as *m* → 0.

Now, considering the ancestral lines that start from two pairs of genes at unlinked loci labeled 1 and 2, the transition matrix from one generation to the previous one takes the form

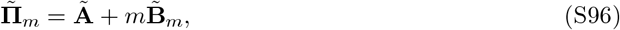

where **Ã** is the transition matrix in the absence of migration. In this case, there will be absorption into 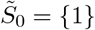, where 1 is the state with only one ancestral line at anyone of the loci, from any initial state with two ancestral lines at each of the loci but at least two in a same deme, which defines the set of states 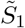. The other states are 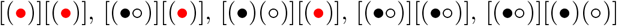 or [(•)(○)] [(•)(○)], where •, ○, and 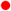 represent sequences ancestral at locus 1 only, at locus 2 only, and at both loci, respectively, while () represents an individual and [] a deme. When they are in this set of states, represented by 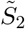, the ancestral lines in each deme will follow a Markov chain on the state space 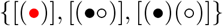, independently of the ancestral lines in the other deme, whose probabilities of transition by recombination or coalescence are given by

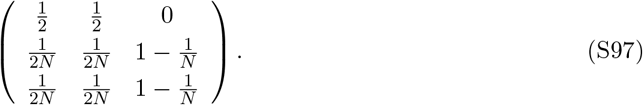

The stationary distribution of this chain is (*π*_1_, *π*_2_, *π*_3_) = (1*/*(*N* + 1), 1*/*(*N* + 1), (*N* − 1)*/*(*N* + 1)). With respect to the state sets 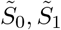 and 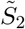 in this order, we have

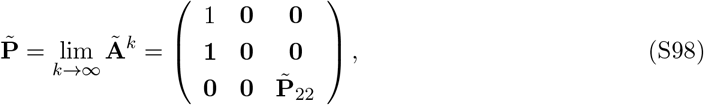

where

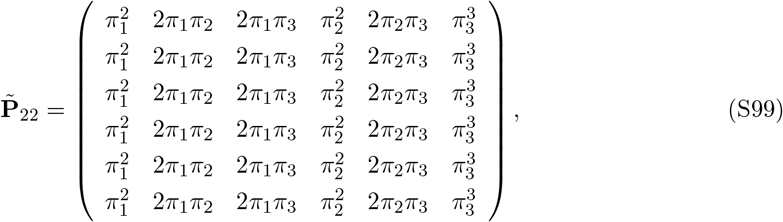

while **0** and **1** are matrices of zeros and ones, respectively. With respect to the same state sets, the rate matrix for changes that involve migration events in the limit as *m* → 0 takes the form

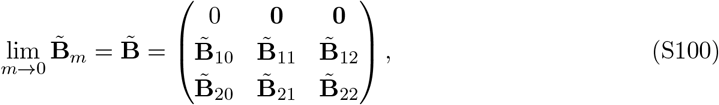

where

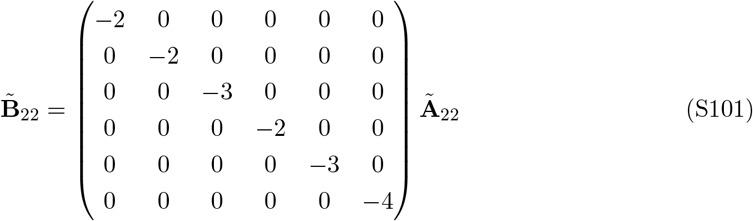

with **Ã**_22_ being the submatrix of **Ã** corresponding to the states in 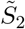 in the above order. Lemma 1 of Möhle (1998a) guarantees that

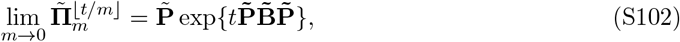

where

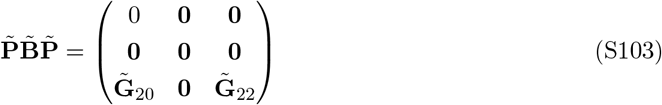

with

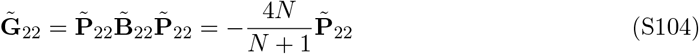

and

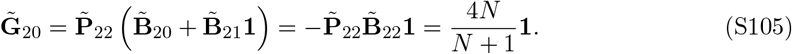

This uses the fact that the entries on each row of 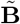 must sum up to 0.

We conclude that, with initial state [(•)][(•)], the survival function satisfies

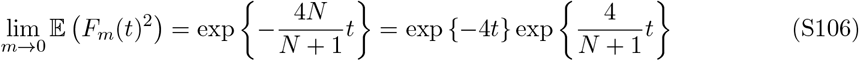

for *t* ≥ 0, which differs slightly from the gametic-migration result (S44). Combining this with our previous result yields

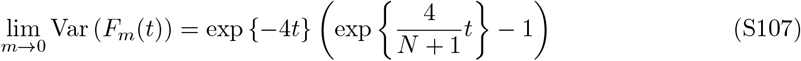

for *t* ≥ 0, which vanishes as *N* → ∞, similar to (S45) under gametic migration.

#### 4.4 Rare-migration limit

With *D* = 2 for fixed *N* and *m* ∈ (0, 1) as *α* → 0, the possible states for the ancestral lines of a pair of genes at a single locus are the same as in the prior subsection and the transition matrix can be expressed as

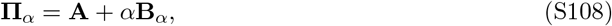

where **A** is given in (S90) and

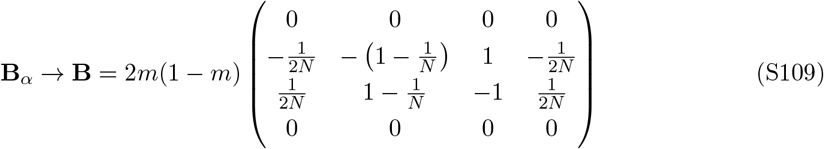

as *α* → 0. Lemma 1 of Möhle (1998a) yields

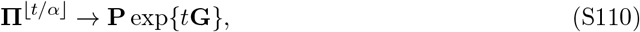

as *α* → 0, where **P** is given in (S93) and

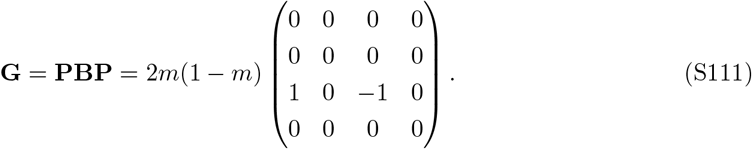

Therefore, for the ancestral lines that start from [(•)][(•)], the survival function satisfies

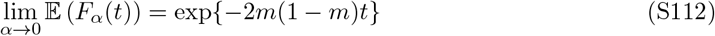

for *t* ≥ 0, which is identical to the corresponding result (S52) for gametic migration. For ancestral lines that start from the initial state [(••)], there is instantaneous coalescence in the limit as *α* → 0.

As for the ancestral lines that start from two pairs of genes at two unlinked loci with the same possible states as in the previous subsection, the transition matrix takes the form

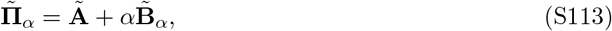

where **Ã** is the same as in the previous subsection, while

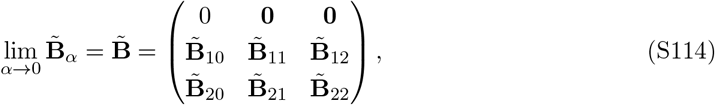

where

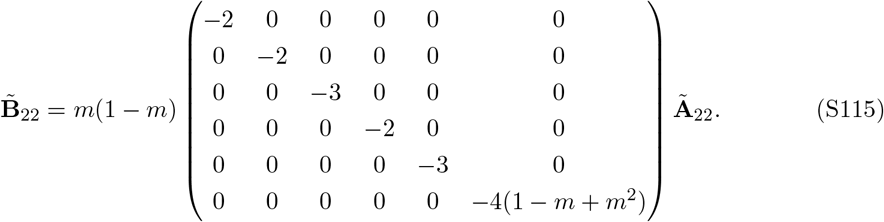

Then Lemma 1 of Möhle (1998a) guarantees that

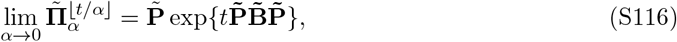

where 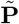 is given in (S98). Then we find

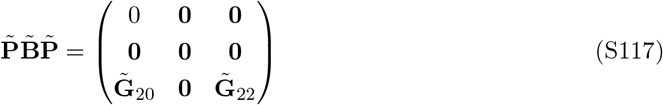

with

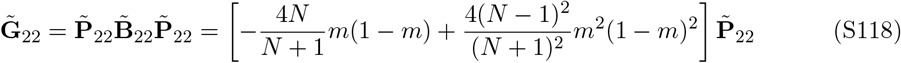

and 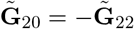. Therefore, with initial state [(•)][(•)], we obtain

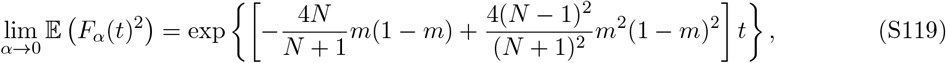

which tends to exp { −4*m*(1 − *m*)(1 − *m* + *m*^2^)*t* } as *N* → ∞, for *t* ≥ 0, and which differs slightly from the gametic-migration result (S63). Overall, although the constants are different, we reach the same conclusion here in the case of diploid migration as we do in the case of gametic migration, namely that the limiting variance of *F*_*α*_(*t*) is not equal to zero in the rare-migration model.

### 5 Dioecious population

Here we assume a dioecious diploid population with *N/*2 males and *N/*2 females in each of *D* demes instaed of a monoecious population. Conditional coalescence times in this model for *D* = 1 were studied in Wakeley et al. (2012), primarily using simulations. For *D* ≥ 2, we will assume either gametic migration as in the main text or diploid migration as in the Section 4. As throughout, *α* is the probability for migration to occur in any given generation. In the case of gametic migration, a proportion *m* of gametes produced in each deme migrate uniformly to the other demes and this is followed by random pairing of male and female gametes within demes. In the case of diploid migration, *m* is the proportion of offspring that uniformly migrate to the other demes after random pairing of male and female gametes within demes.

The state space for two distinct ancestral lines at the same locus is the same as in Section 4 for diploid migration in a monoecious population: [(•)] for one in one individual in one deme, [(•)(•)] for two in two individuals in the same deme, [(•)][(•)] for two in two individuals in different demes, and [(••)] for two in the same individual in one deme. The transition matrix is slightly different, being given by

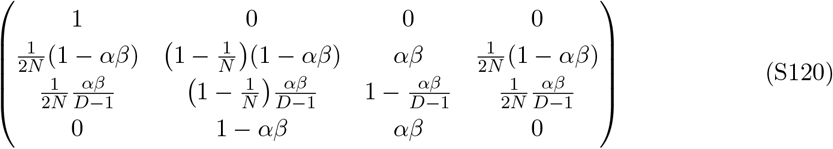

under gametic migration and

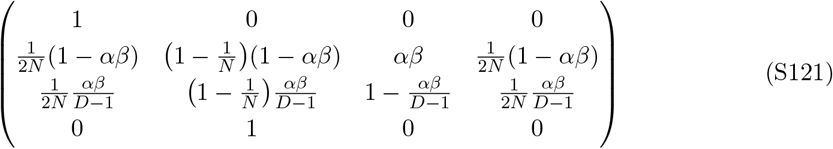

under diploid migration. Here,

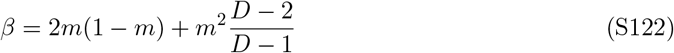

represents the conditional probability for two gametes or offspring, respectively, chosen at random in the same deme to come from two different demes given that migration occurs, and *β/*(*D* − 1) the conditional probability for two gametes or offspring, respectively, in different demes to come from the same deme.

#### 5.1 Structured-coalescent limit

For the structured-coalescent with *D* = 2, *α* = 1 and *m* = *M/*(4*N*), the transition matrix for a pair of ancestral lines at a single locus can be written in the form

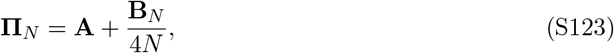

where **A** is the same as in Subsection 4.1, while **B** = lim_*N*→∞_ **B**_*N*_ is given by

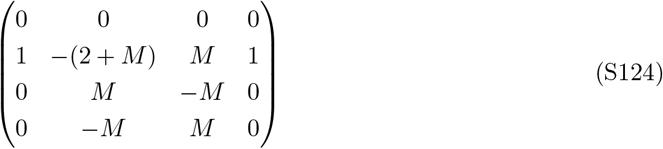

under gametic migration and

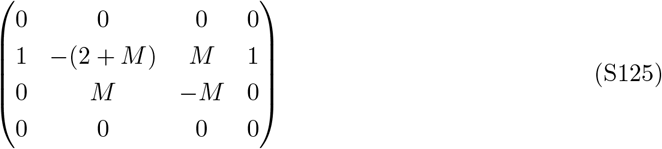

under diploid migration. In both cases, **PBP** where **P** = lim_*k*→∞_ **A**^*k*^ = **A** is the same as in Subsection 4.1.

For two pairs of ancestral lines at two unlinked loci, the transition matrix takes the form

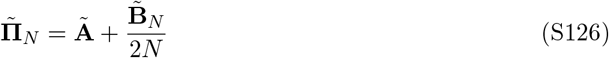

with the same 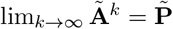 and the same first 6 rows of 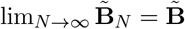 corresponding to the states with fewer than four ancestral lines or four ancestral lines in four different individuals. The same rate matrix 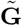 as in (S76) and (22) is obtained, and the same conclusion that the effect of the pedigree becomes negligible ensues because lim_*N*→∞_ Var(*F*_*N*_ (*t*)) = 0.

#### 5.2 Many-demes limit

With *D* ≥ 4 and *α* = 1, the transition matrix for a pair of ancestral lines at the same locus can be decomposed as

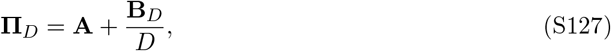

where

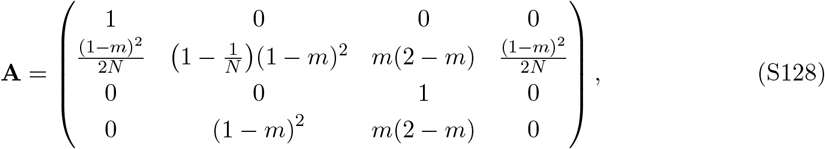

under gametic migration and

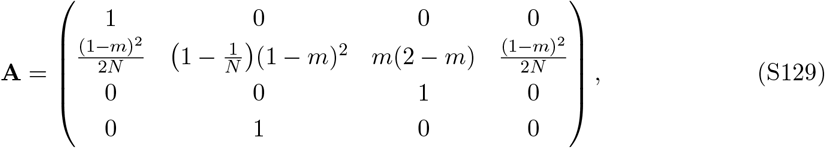

under diploid migration. In both cases, we have

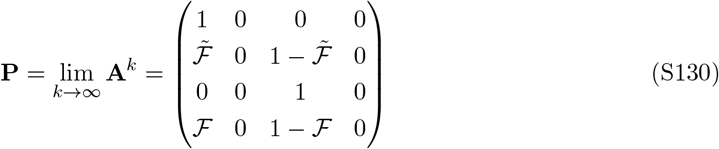

as in Subsection 4.2, but where

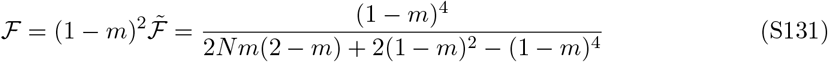

under gametic migration and

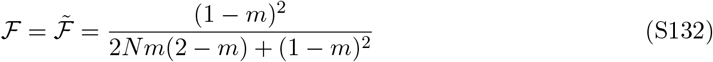

under diploid migration. On the other hand,

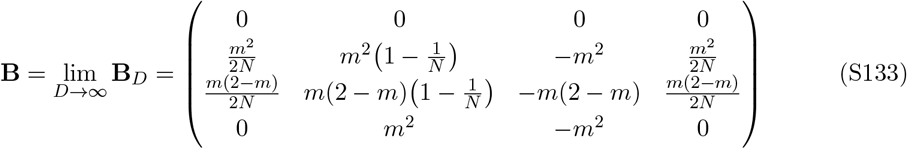

under gametic migration, while it is the same as in Subsection 4.2 with all zero entries in the last row under diploid migration. This leads to the rate matrix

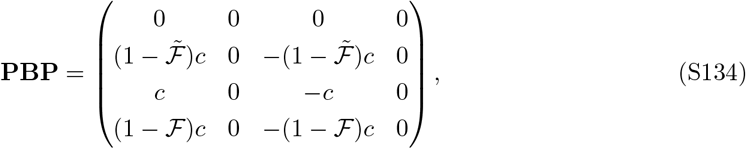

where

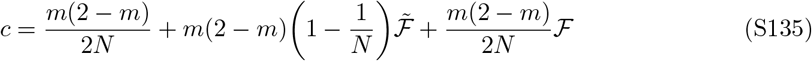

is the rate of coalescence starting from state [(•)][(•)] as in Subsection 4.2. Here, we have

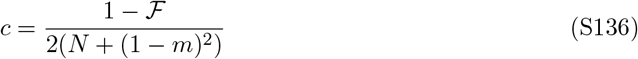

under gametic migration and

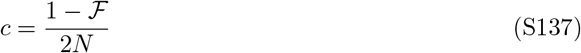

under diploid migration. This means that there is instantaneous coalescence from states [(•)(•)] and [(••)] with probabilities 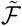 and ℱ, respectively, and instantaneous transition to state [(•)][(•)] with the complementary probabilities from which coalescence occurs at rate *c*.

Similarly, starting from two ancestral lines at each of two unlinked loci in four individuals in different demes, the rate of coalescence is

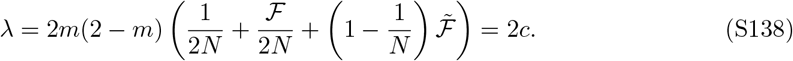

On the other hand, starting from two ancestral lines at each of two unlinked loci in the same individual in the same deme, there will be either instantaneous coalescence with some probability 𝒰 or instantaneous transition to all ancestral lines in different individuals in different demes with probability 1 − 𝒰, so that lim_*D*→∞_ Var(*F*_*D*_(*t*)) ≠ 0 in general. But in the limit of a large deme size with *M* = 4*Nm* kept constant, we have

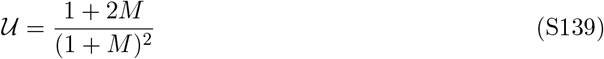

as in Subsection 4.2, so that the effect of the pedigree does become negligible with the addition of this structured-coalescent assumption.

#### 5.3 Low-migration limit

With *D* = 2, *α* = 1 and considering a small migration fraction *m*, the transition matrix for a pair of ancestral lines at a single locus takes the form

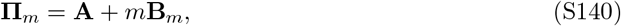

where

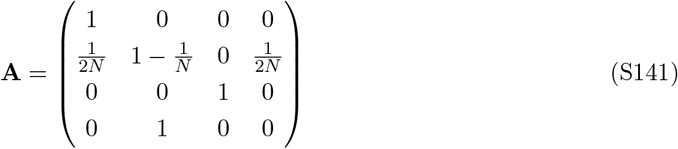

under either gametic migration or diploid migration, while **B** = lim_*m*→0_ **B**_*m*_ is given by

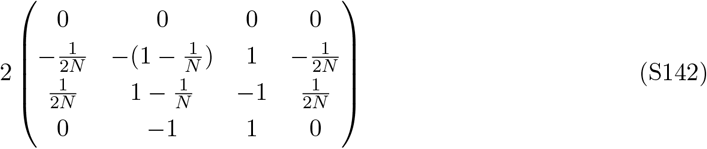

under gametic migration and the same matrix but with all zero entries in the last row as in Subsection 4.3 under diploid migration. This leads to the same **P** = lim_*k*→∞_ **A**^*k*^ and the rate matrix **PBP** in the limit *m* → 0 with time measured in units of 1*/m* generations. This means instant coalescence for two ancestral lines at the same locus if they start from the same deme, but coalescence at rate 2 if they start from different demes.

Now, considering the ancestral lines that start from two pairs of genes at two unlinked loci and arranging the states as in Subsection 4.3, the transition matrix from one generation to the previous one takes the same form

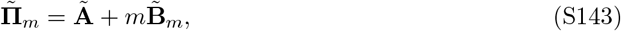

where

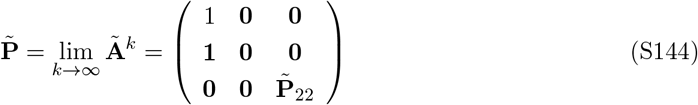

with each row of 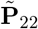 given by 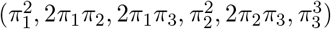. Here, the frequency vector (*π*_1_, *π*_2_, *π*_3_) = (1*/*(*N* + 2), 1*/*(*N* + 2), *N/*(*N* + 2)) is the stationary distribution of the transition matrix

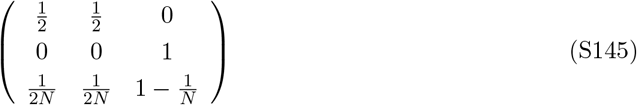

for two ancestral lines at two unlinked loci on the same chromosome, on different chromosomes in the same individuals and in different individuals, respectively, in the same deme in the absence of migration. On the other hand,

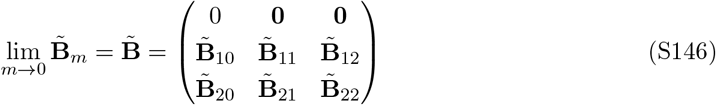

has

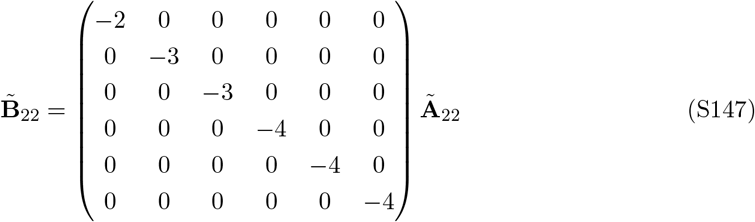

under gametic migration and the same 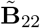 as in Subsection 4.3 under diploid migration. Starting from two ancestral lines at each of two loci in different demes, this gives a rate of coalescence

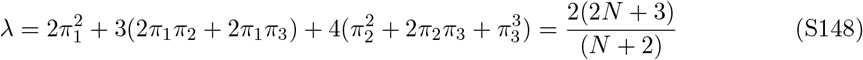

under gametic migration and

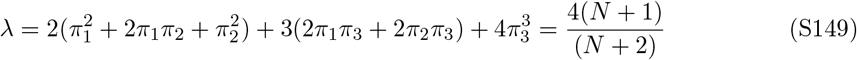

under diploid migration. Otherwise, there is instantaneous coalescence. Notice that *λ* → 4 as *N* → ∞. This is the same rate as in the diploid, monoecious migration model of Subsection 4.3, and is twice the rate for the haploid migration model in Section 2. Overall, the same conclusions hold, that there is an effect of the pedigree but that it becomes negligible as the deme size increases.

#### 5.4 Rare-migration limit

In the case of a fixed migration fraction *m* ∈ (0, 1) with small probability *α* in each generation for *D* = 2, the transition matrix for a pair of ancestral lines at a single locus takes the form

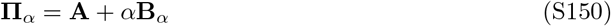

with the same **A** as in the previous subsection but with a multiplicative factor 2*m*(1 − *m*) instead of 2 in **B** = lim_*α*→0_ **B**_*α*_, which gives the rate of coalescence of two ancestral lines starting from different demes. This is the same constant 2*m*(1 − *m*) that appears in Section 3 and Subsection 4.4.

For two pairs of ancestral lines at two unlinked loci, the decomposition

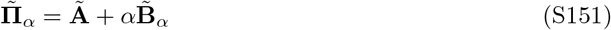

has the same **Ã** as in the previous subsection, while 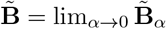 has

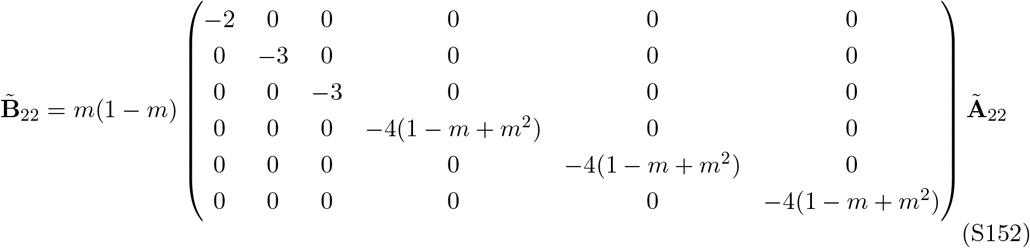

under gametic migration and the same 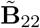 as in Subsection 4.4 under diploid migration and monoecy. Therefore, there is instantaneous coalescence unless the two ancestral lines at each of the two loci are in different demes, in which case the rate of coalescence is given by

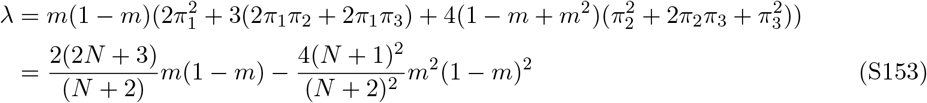

under gametic migration and

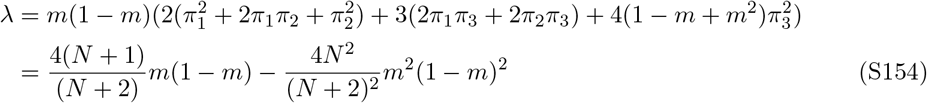

under diploid migration. In both cases, *λ* → 4*m*(1 − *m*)(1 − *m* + *m*^2^) as *N* → ∞. Again, although the constants differ, the conclusion is the same as in Rare-migration limit under gametic migration, that there is a persistent pedigree effect for *m* ∈ (0, 1).

### 6 Summary table of results

Our results on the limiting variance of the conditional survival function for all the models are summarized in Table S1 where

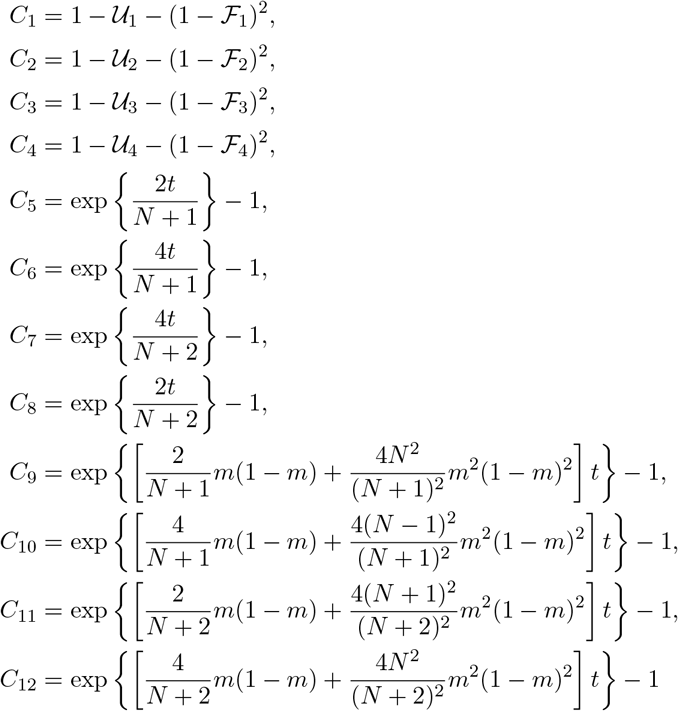

with

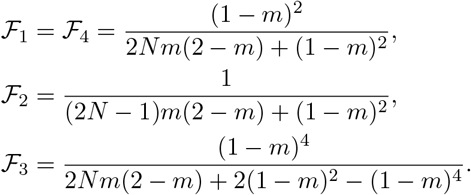

Here, 𝒰_1_, 𝒰_2_, 𝒰_3_, 𝒰_4_ are probabilities of coalescence at any of two unlinked loci for ancestral lines starting in the same individual in the limit of a large number of demes, while ℱ_1_, ℱ_2_, ℱ_3_, ℱ_4_ are probabilities of coalescence for ancestral lines at a single locus under the same assumptions. With *M* = 4*Mn* held fixed as *N* → ∞, the former have the same limit (1 + 2*M*)*/*(1 + *M*)^2^, and the latter the same limit 1*/*(1 + *M*). Moreover, *C*_1_, …, *C*_8_ tend to 0 as *N* → ∞ and this occurs when ancestral lines at different loci cannot stay on the same chromosomes in the limit of a large deme size so that coalescence times at unlinked loci are independent in this limit. This is not the case in the rare-migration limit, however, with *C*_9_, …, *C*_12_ all tending to exp {[4*m*^2^(1 − *m*)^2^ *t*} − 1 *>* 0.

**Table S1:**
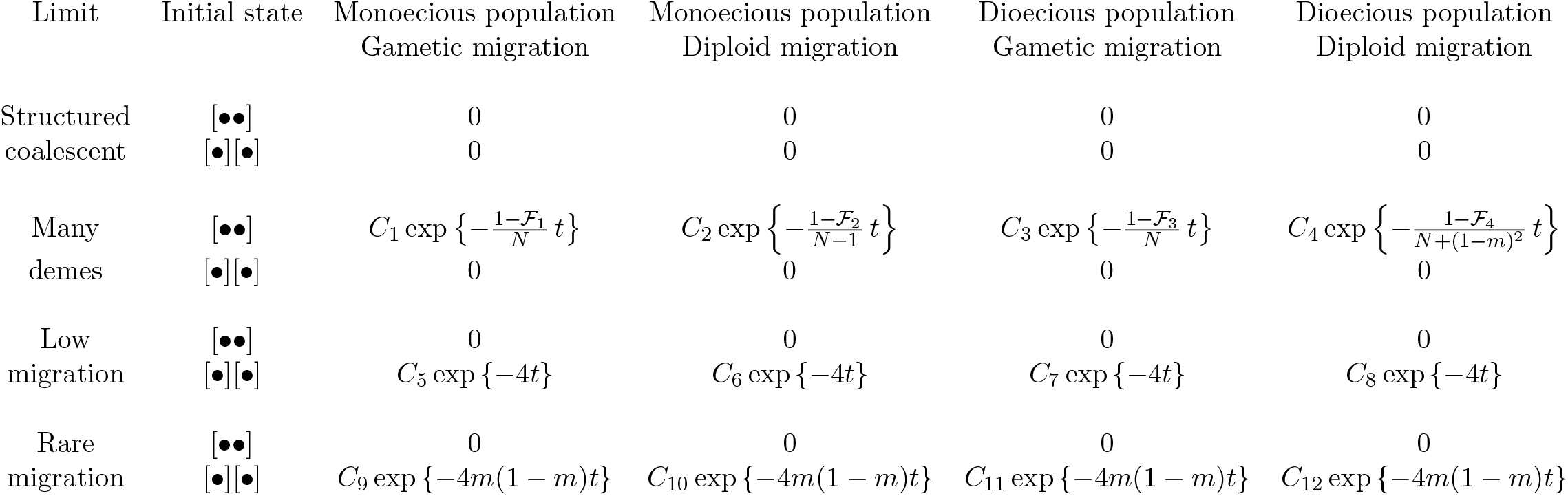
Limiting variance of the conditional survival function given the population pedigree. The exponential functions are the squares of the limiting expectations. In the low- and rare-migration limits with initial state [••], the survival function takes the value 0 with probabilty 1. With diploid migration, the initial states [••] and [•][•] are replaced by [(••)] and [(•)][(•)], respectively.

