## Supplementary material for "A classification of structured coalescent processes with migration, conditional on the population pedigree": associated_code: low_rare_sim_plot.jl

```
using JLD2, FileIO, Statistics, Plots, Random, Measures, LaTeXStrings
gr()

# -----------------------------
# Global style
# -----------------------------
default(fontfamily="Helvetica", framestyle=:box)

label_fs  = 14
legend_fs = 14

# ---- OKABE–ITO PALETTE ----
oi_skyblue = RGB(0.34, 0.71, 0.91)
oi_bluishg = RGB(0.00, 0.62, 0.45)
oi_verm    = RGB(0.84, 0.37, 0.00)
oi_purple  = RGB(0.80, 0.47, 0.65)

# ---- FINAL COLOR CHOICES (YOUR REQUEST) ----
clr_spaghetti = RGB(0.12, 0.47, 0.71)   # blue spaghetti
clr_mean      = oi_verm                 # Okabe–Ito orange
clr_theory    = RGB(0,0,0)              # BLACK theory

# -----------------------------
# Load data
# -----------------------------
data = <inserst jld2 file path here>

N_val     = data["dN"][1]
desired_m = data["m_values"][1]
p_mig     = 0.001  # migration event probability (α)

# Extract coalescence time vectors
ecdf_data_list = [data["all_results"][run][N_val][desired_m]
                  for run in 1:data["num_runs"]]
total_runs = length(ecdf_data_list)
println("Total simulation runs: ", total_runs)

# Random selection of CDFs for plotting
num_random_cdfs = min(300, total_runs)
println("Randomly selecting ", num_random_cdfs, " CDFs from ", total_runs)
random_indices = shuffle(1:total_runs)[1:num_random_cdfs]

# -----------------------------
# ECDF grid + rescaling (keep your original scaling)
# -----------------------------
min_val = minimum(minimum.(ecdf_data_list))
max_val = maximum(maximum.(ecdf_data_list))

grid        = collect(range(min_val, max_val, length=200))
grid_scaled = grid .* p_mig   # time × α

function evaluate_ecdf(sorted_data::Vector{<:Real}, grid::AbstractVector{<:Real})
    n = length(sorted_data)
    return [searchsortedlast(sorted_data, t) / n for t in grid]
end

# Build ECDF matrix
cdf_matrix = zeros(total_runs, length(grid))
for i in 1:total_runs
    sorted_data = sort(ecdf_data_list[i])
    cdf_matrix[i, :] = evaluate_ecdf(sorted_data, grid)
end

mean_ecdf = vec(mean(cdf_matrix, dims=1))

# -----------------------------
# Theory curve
# -----------------------------
gamma = 2 * desired_m * (1 - desired_m) #change to 2 for low-migration
theoretical_curve = 1 .- exp.(-gamma .* grid_scaled)

# -----------------------------
# Truncate when CDF values approach 1 to avoid long right tail
# -----------------------------
cutoff_value = 0.998
cutoff_index = findfirst(x -> x >= cutoff_value, mean_ecdf)
cutoff_index = cutoff_index === nothing ? length(grid) : cutoff_index

trimmed_grid_scaled       = grid_scaled[1:cutoff_index]
trimmed_cdf_matrix        = cdf_matrix[:, 1:cutoff_index]
trimmed_mean_ecdf         = mean_ecdf[1:cutoff_index]
trimmed_theoretical_curve = theoretical_curve[1:cutoff_index]

println("Trimming time range at index $(cutoff_index), mean ECDF ≈ $(trimmed_mean_ecdf[end])")

# -----------------------------
# Plot
# -----------------------------
main_title = "Rare -migration (m = $(desired_m), alpha = $(p_mig))"  #change for low-migration

plt = plot(
    xlabel="Time in units of 1/alpha generations", #change for low-migration 1/m
    ylabel="CDF",
    title=main_title,
    titlefont=font(label_fs, "Helvetica"),
    size=(720, 540),
    margin=6mm,
    grid=false,
    legend=:bottomright,
    legend_background_color=:transparent,
    legend_foreground_color=:transparent,
    guidefontsize=label_fs,
    tickfontsize=label_fs,
    legendfontsize=legend_fs,
    ylims=(0, 1.01)
)

# ---- SPAGHETTI ECDFs (BLUE) ----
for k in 1:num_random_cdfs
    idx = random_indices[k]
    plot!(plt,
          trimmed_grid_scaled,
          trimmed_cdf_matrix[idx, :];
          lw=1.0,
          color=clr_spaghetti,
          alpha=0.45,
          label=""
    )
end

plot!(plt, [NaN], [NaN];
      lw=2.0,
      color=clr_spaghetti,
      alpha=1.0,
      label="Emprical Conditional CDF"
)

# ---- Mean ECDF (OKABE–ITO ORANGE) ----
plot!(plt,
      trimmed_grid_scaled,
      trimmed_mean_ecdf;
      lw=2.8,
      color=clr_mean,
      label="Emprical Average CDF"
)

# ---- Theory (BLACK, dashed) ----
plot!(plt,
      trimmed_grid_scaled,
      trimmed_theoretical_curve;
      lw=2.6,
      linestyle=:dash,
      color=clr_theory,
      label="Marginal CDF"
)

display(plt)
```
