## Supplementary material for "A classification of structured coalescent processes with migration, conditional on the population pedigree": associated_code: many_demes_leading_coeff_decay.jl

```
using Statistics, Plots, Measures
gr()
default(fontfamily="Helvetica", framestyle=:box)


# Parameters

Mmin     = 1e-2
Mmax     = 1e3
grid_len = 800
label_fs = 14


# Okabe–Ito colour colourblind friendly

oi_verm = RGB(0.84, 0.37, 0.00)
oi_bluishg = RGB(0.00, 0.62, 0.45)

from Supplementary 
# g(M) = lim_{N->∞, 4Nm=M} N*(1 - u17 - (1 - F)^2)

g(M) = M*(M^2 + 4M + 6) / (3*(1 + M)^3)

# Log10-spaced grid in M
Mgrid = exp10.(range(log10(Mmin), log10(Mmax), length=grid_len))
gvals = g.(Mgrid)

# Custom ticks (PDF-safe)
xtick_vals  = [1e-2, 1e-1, 1e0, 1e1, 1e2, 1e3]
xtick_labels = ["10e-2","10e-1","10e0","10e1","10e2","10e3"]


# Plot 

plt = plot(
    Mgrid, gvals;
    xlabel="4Nm = M (log scale)",
    ylabel="g(M)",
    title="Leading-order Coefficient in Decay of Variance ",
    titlefont=font(label_fs, "Helvetica", :bold),
    size=(720, 540),
    margin=6mm,
    grid=false,
    legend=false,
    guidefontsize=label_fs,
    tickfontsize=label_fs,
    lw=3,
    color=oi_bluishg,
    xscale=:log10,
    xticks=(xtick_vals, xtick_labels),
    xlims=(Mmin, Mmax)
)

display(plt)
```
