## Supplementary material for "A classification of structured coalescent processes with migration, conditional on the population pedigree": associated_code: simulate_coalescence.jl

```
#=
Packages

Random:
  Used for all stochastic elements of the simulation, including
  pedigree construction and coalescence.

JLD2:
  Used to save simulation output and parameters to disk in .jld2 format
  for later analysis and plotting.

FileIO:
  Included for JLD2/FileIO compatibility when saving/loading .jld2 files.
=#

using Random
using JLD2
using FileIO


#=
Pedigree generator for a diploid Wright–Fisher population with two demes
and gametic migration (D = 2).

Time:
  g counts generations backward from the sampled present.
  g = 0 is present; parents are at generation g+1.

Migration model:
  - α : probability that migration is possible in a given generation
  - m        : per-gamete migration probability, conditional on migration
               being possible in that generation

If migration is not possible in a generation (with probability 1−α),
all gametes remain in their parental deme.
=#

function generate_pedigree(N::Int, T_max::Int, m::Float64, α::Float64; D::Int=2)

    # This generator is currently specialized to D = 2.
    
    # ped stores the pedigree:
    #
    #   key   = (g, d, i)
    #           g : generation index (backward from present)
    #           d : deme (1 or 2)
    #           i : individual index within deme
    #
    #   value = ((pd1, p1), (pd2, p2))
    #           (pd1, p1) : deme and index of first parental gamete
    #           (pd2, p2) : deme and index of second parental gamete
    #
    # Parents are in the previous generation g+1.
    #
    ped = Dict{Tuple{Int,Int,Int}, Tuple{Tuple{Int,Int},Tuple{Int,Int}}}()

    # Helper for switching demes under migration
    other_deme(d) = (d == 1 ? 2 : 1)

    # Loop over generations
    for g in 0:(T_max-1)

        # Decide whether migration is possible this generation
        mig_on = rand() < α

        # Loop over demes and individuals
        for d in 1:D, i in 1:N

            # Each gamete independently migrates with probability m,
            # conditional on migration being possible this generation
            p1 = ((mig_on && rand() < m) ? other_deme(d) : d, rand(1:N))
            p2 = ((mig_on && rand() < m) ? other_deme(d) : d, rand(1:N))

            # Record the two parental gametes for individual (g,d,i)
            ped[(g,d,i)] = (p1, p2)
        end
    end

    # Return the full pedigree
    return ped
end


#=
Simulate the ancestry of two gene copies on a fixed pedigree
and record the time to coalescence (in generations back).

State = (generation g, deme d, individual i, gene a)
  a ∈ {1,2} denotes which gene copy inside the individual.

Conventions:
  - If both lineages hit the same (d,i) with the same gene a, they have coalesced.
  - If they hit the same (d,i) but different genes, they must move to different parental
    gametes in the next generation (no independent choices that could keep them identical).

Return:
  The first generation t where coalescence occurs (t ≥ 0),
  measured in generations backward from the present.
=#

function simulate_coalescence(ped, N::Int, T_max::Int; deme::Int=1)

    # ---- initial configuration (present generation, g = 0) ----
    #
    # Sample two different individuals in the same sampled deme.
    #
    i1 = rand(1:N)
    i2 = rand(1:N)
    while i2 == i1
        i2 = rand(1:N)
    end

    # Sample one gene copy from each individual at time 0
    a1 = rand(Bool) ? 1 : 2
    a2 = rand(Bool) ? 1 : 2

    s1 = (0, deme, i1, a1)
    s2 = (0, deme, i2, a2)

    # ---- trace ancestry backward generation by generation ----
    for t in 1:T_max

        g1, d1, i1, a1 = s1
        g2, d2, i2, a2 = s2

        # Get the two parental gametes for each current individual
        p1 = ped[(g1, d1, i1)]
        p2 = ped[(g2, d2, i2)]

        # If both lineages are already in the same individual:
        if d1 == d2 && i1 == i2

            # Same gene => already identical before stepping this generation
            if a1 == a2
                return t - 1
            else
                # Different genes => must follow different parental gametes
                # gene 1 follows parental gamete 1; gene 2 follows parental gamete 2
                n1_d, n1_i = p1[a1]
                n2_d, n2_i = p2[a2]

                # After moving into the parental individual, choose a random gene there
                s1 = (g1 + 1, n1_d, n1_i, rand(Bool) ? 1 : 2)
                s2 = (g2 + 1, n2_d, n2_i, rand(Bool) ? 1 : 2)
                continue
            end
        end

        # Otherwise: different individuals (or different demes)
        # Each lineage chooses which parental gamete carries the locus (50/50)
        link1 = rand(Bool) ? 1 : 2
        link2 = rand(Bool) ? 1 : 2

        n1_d, n1_i = p1[link1]
        n2_d, n2_i = p2[link2]

        # Move one generation back and choose a random gene in the parent
        s1 = (g1 + 1, n1_d, n1_i, rand(Bool) ? 1 : 2)
        s2 = (g2 + 1, n2_d, n2_i, rand(Bool) ? 1 : 2)
    end

    # If coalescence was not observed before the cutoff, return T_max
    return T_max
end


#=
Simulation parameters (structured-coalescent regime)
=#

# N : number of diploid individuals per deme
N = 124

# D : number of demes (this script assumes D = 2)
D = 2

# M : scaled migration rate (structured-coalescent parameterization)
M = 0.5

# α : probability that migration is possible in a given generation
α = 1.0

# m : per-gamete migration probability when migration is possible
m = M / (4N)

# T_max : maximum number of generations traced backward
T_max = 50_000

# num_ped : number of independent pedigrees ("runs" in the saved file)
num_ped = 1000

# num_rep : number of genealogies (loci) simulated per pedigree
num_rep = 10_000

# Random seed for reproducibility
Random.seed!(1)


#=
Set the name of the output file here.
Use something descriptive (regime, N, M, seed, etc.).
=#

out_file = <insert jld2 file name>


#=
Run simulations

Generates num_ped independent pedigrees. For each pedigree, simulates
num_rep coalescence times conditional on that pedigree.

Results are stored in the schema the plotting/loader expects:
  all_results[r][N][m_key] = Vector{Int} of coalescence times (generations)
=#

all_results = Vector{Dict{Int, Dict{Float64, Vector{Int}}}}(undef, num_ped)

m_key = Float64(m)

for r in 1:num_ped

    # Generate one pedigree
    ped = generate_pedigree(N, T_max, m, α; D=D)

    # Simulate genealogies conditional on this pedigree
    times_gen = Vector{Int}(undef, num_rep)
    for k in 1:num_rep
        times_gen[k] = simulate_coalescence(ped, N, T_max; deme=1)
    end

    # Store results in loader-compatible structure
    all_results[r] = Dict(N => Dict(m_key => times_gen))

    # Progress message
    r % 10 == 0 && println("Completed pedigree $r / $num_ped")
end


#=
Save results

The following quantities are saved:

  all_results : simulation output (one dict per pedigree)
  dN          : vector containing N
  m_values    : vector containing m_key
  num_runs    : number of independent pedigrees (num_ped)
  α           : probability migration is possible in a generation
=#

dN = [N]
m_values = [m_key]
num_runs = num_ped

JLD2.@save out_file dN m_values num_runs α all_results

println("Saved to: ", out_file)
```
