## Supplementary material for "A classification of structured coalescent processes with migration, conditional on the population pedigree": associated_code: structcoal_sim_plot.jl

```
#=
Packages

JLD2:
  Used to load simulation output saved in .jld2 format.

Plots:
  Used to render plots

Measures:
  Used for plot margins (e.g., 6mm).
=#

using JLD2
using Statistics
using Plots
using Measures
using Random
using LinearAlgebra

gr()

default(fontfamily="Helvetica", framestyle=:box)

label_font_size  = 14
legend_font_size = 14

# Okabe–Ito palette (RGB)
oi_vermilion = RGB(0.84, 0.37, 0.00)
oi_blue      = RGB(0.00, 0.45, 0.70)

# Plot colours
colour_first  = RGB(0.0, 0.62, 0.45)   # plotted FIRST
colour_second = oi_vermilion           # plotted SECOND (spaghetti on top)
colour_third  = oi_blue                # plotted THIRD  (spaghetti on top)

grid_len   = 300
cdf_cutoff = 0.999

# Plot only up to this time (x-axis) in units of 2N generations
tmax_2N = 15.0

# Number of spaghetti curves to show (conditional CDFs)
n_spaghetti = 1000

# Migration rate for theoretical curve (must match the M used in your model, in coalescent units)
M_theory = 0.5

file_first  = <insertjld2 file name > 
file_second = <insertjld2 file name > 
file_third  = <insertjld2 file name > 


# Marginal Structured Coalescent CDF

# CDF:
#   F(t) = P(T ≤ t) = 1 - P(T > t)
#
function cdf_structured_coalescent_state2(t::Real, M::Real, state)
    @assert M > 0 "M must be positive"

    G = [
        0.0   0.0        0.0
        1.0  -(1.0 + M)  M
        0.0   M         -M
    ]

    v = state #[0.0, 1.0, 0.0]   # start in state 2 (same deme)
    # v = [0.0, 0.0, 1.0] # uncomment to start in state 3 (different demes)
    w = [0.0, 1.0, 1.0]   # survival = being in states 2 or 3

    survival = (v' * exp(t * G) * w)[1]
    return 1 - survival
end


# CDF evaluation on a grid

function evaluate_ecdf(sorted_samples::AbstractVector{<:Real},
                       grid::AbstractVector{<:Real})
    n = length(sorted_samples)
    return [searchsortedlast(sorted_samples, t) / n for t in grid]
end

#=
Load + process one file
=#
function process_file_spaghetti_mean(results_file::AbstractString; grid_len::Int=300, n_spaghetti::Int=300)
    d = JLD2.load(results_file)

    N     = d["dN"][1]
    m_key = d["m_values"][1]
    R     = d["num_runs"]

    runs = [d["all_results"][run][N][m_key] for run in 1:R]

    min_time = minimum(minimum.(runs))
    max_time = maximum(maximum.(runs))
    ecdf_grid = collect(range(min_time, max_time, length=grid_len))

    # Convert to units of 2N generations on x-axis
    grid_scaled = ecdf_grid ./ (2N)

    Fmat = zeros(length(runs), length(ecdf_grid))
    for i in eachindex(runs)
        sorted_times = sort(runs[i])
        Fmat[i, :] .= evaluate_ecdf(sorted_times, ecdf_grid)
    end

    mean_ecdf = vec(mean(Fmat, dims=1))

    k = min(n_spaghetti, size(Fmat, 1))
    idx = randperm(size(Fmat, 1))[1:k]
    spaghetti = Fmat[idx, :]

    return (N=N, m_key=m_key, grid=grid_scaled, spaghetti=spaghetti, mean=mean_ecdf)
end

r1 = process_file_spaghetti_mean(file_first;  grid_len=grid_len, n_spaghetti=n_spaghetti)
r2 = process_file_spaghetti_mean(file_second; grid_len=grid_len, n_spaghetti=n_spaghetti)
r3 = process_file_spaghetti_mean(file_third;  grid_len=grid_len, n_spaghetti=n_spaghetti)

function trim_at_time(r, tmax::Real)
    idx = findlast(x -> x <= tmax, r.grid)
    idx = idx === nothing ? 1 : idx
    return (N=r.N, m_key=r.m_key,
            grid=r.grid[1:idx],
            spaghetti=r.spaghetti[:, 1:idx],
            mean=r.mean[1:idx],
            tmax_idx=idx)
end

r1_t = trim_at_time(r1, tmax_2N)
r2_t = trim_at_time(r2, tmax_2N)
r3_t = trim_at_time(r3, tmax_2N)


# Theoretical curve on same x-axis
theory_grid = r1_t.grid
theory_cdf  = [cdf_structured_coalescent_state2(τ, M_theory, [0.0,1.0,0.0]) for τ in theory_grid] #same deme
theory_cdf2  = [cdf_structured_coalescent_state2(τ, M_theory, [0.0,0.0,1.0]) for τ in theory_grid] #different deme

# Plot

plt = plot(
    xlabel="Time in units of 2N generations",
    ylabel="CDF",
    title="Structured -Coalescent Approximation for M=0.5",
    titlefont=font(label_font_size, "Helvetica"),
    size=(720, 540),
    margin=6mm,
    grid=false,
    legend=:bottomright,
    legend_background_colour=:transparent,
    legend_foreground_colour=:transparent,
    guidefontsize=label_font_size,
    tickfontsize=label_font_size,
    legendfontsize=legend_font_size,
    ylims=(0, 1.01),
    xlims=(0, tmax_2N)
)

# spaghetti
sp_lw = 1.0
sp_alpha_first = 0.12
sp_alpha_top   = 0.25

for i in 1:size(r1_t.spaghetti, 1)
    plot!(plt, r1_t.grid, vec(r1_t.spaghetti[i, :]);
          lw=sp_lw, color=colour_first, alpha=sp_alpha_first, label="")
end

for i in 1:size(r2_t.spaghetti, 1)
    plot!(plt, r2_t.grid, vec(r2_t.spaghetti[i, :]);
          lw=sp_lw, color=colour_second, alpha=sp_alpha_top, label="")
end

for i in 1:size(r3_t.spaghetti, 1)
    plot!(plt, r3_t.grid, vec(r3_t.spaghetti[i, :]);
          lw=sp_lw, color=:black, alpha=sp_alpha_top, label="")
end

# Legend 
plot!(plt, [NaN], [NaN]; lw=2.5, color=colour_first,  alpha=1.0, label="Empirical Conditional CDF N=24")
plot!(plt, [NaN], [NaN]; lw=2.5, color=colour_second, alpha=1.0, label="Empirical Conditional CDF N=124")
plot!(plt, [NaN], [NaN]; lw=2.5, color=:black,        alpha=1.0, label="Empirical Conditional CDF N=640")

# Emprical Average
plot!(plt, r1_t.grid, r1_t.mean;lw=2.7, color=colour_first, linestyle=:solid, label="Emprical Average CDF N=24")

# Marginal CDF in red
plot!(plt, theory_grid, theory_cdf; lw=2.7, color="#F0E442", linestyle=:solid, label="Marginal CDF")
#plot!(plt, theory_grid, theory_cdf2; lw=2.7, color=:red, linestyle=:solid, label="Marginal CDF")


display(plt)
```
